# Environmental and mutational modulation of collateral fitness effects informs their mechanisms

**DOI:** 10.64898/2026.01.22.699087

**Authors:** Cameron Goff, Erh-Yeh Tsou, Jacob D. Mehlhoff, Marc Ostermeier

## Abstract

Fitness effects of mutations that do not arise from changes in a protein’s ability to perform its physiological functions (called collateral fitness effects or CFEs) are an understudied aspect of fitness landscapes. We have previously systematically measured the CFEs of all possible single amino acid substitutions in four proteins and found the frequency of deleterious mutations to vary by two orders of magnitude. Of these proteins, TEM-1 β-lactamase had the highest frequency, and deleterious mutations caused TEM-1 aggregation. Here, we systematically measured TEM-1 collateral fitness landscapes in environments and situations expected to alter protein aggregation or protein stability. We found a moderate correlation between deleterious CFEs and predicted thermodynamic stability effects in TEM-1’s α-domain. Empirically, we found that the frequency and magnitude of deleterious CFEs can be reduced by altering the growth environment to disfavor aggregation (i.e. reducing the growth temperature or shifting to minimal media) or by stabilizing TEM-1 (via the M182T mutation or the addition of the β-lactamase inhibitor avibactam to the growth medium). However, although raising the growth temperature to favor aggregation exacerbated deleterious CFEs of many mutations, many mutations’ effects were reduced. Furthermore, although reductions in CFEs occurred with reductions in TEM-1 aggregation for some mutants, for many mutants they did not. We propose that mutational destabilization exposes protein motifs that can cause deleterious CFEs, but that these motifs and those that cause aggregation are not necessarily the same motifs.

## Introduction

We have previously proposed that fitness effects of protein mutations can be classified into two broad categories (Mehlhoff, et al. 2020). Primary fitness effects (PFEs) are those that arise from changes in the ability of the protein to perform its physiological function. Collateral fitness effects (CFEs) are those that do not arise from changes in the ability of the protein to perform its physiological function. Potential examples of CFEs include mutations leading to the accumulation of toxic protein aggregates and mutations that cause new and injurious interactions with other biomolecules (e.g., inhibition of an essential enzyme or interference with protein export across membranes). The distinction between PFEs and CFEs is important for several reasons. First, CFEs have been underappreciated as a source of constraints on protein evolution. Second, CFEs constrain evolution whenever the protein is expressed whereas PFEs constrain evolution only when the protein’s physiological function is relevant to fitness. Third, CFEs are a possible contributing factor to the phenomena of the E-R anticorrelation (Pal, et al. 2001; Zhang and Yang 2015), the observation that the best predictor of protein evolution rates is protein expression level, with more highly expressed proteins tending to evolve more slowly. Fourth, several diseases are associated with protein misfolding, though the study of the source of the relationship and whether misfolding is disease causative has often been vexing. The study of the cause of CFEs might be informative in this regard.

Previously, we systematically and comprehensively measured the CFEs of single amino acid substitutions in four antibiotic resistance proteins: TEM-1 β-lactamase, New Delhi metallo-β-lactamase (NDM-1), chloramphenicol acetyltransferase I (CAT-I), and 2″-aminoglycoside nucleotidyltransferase (AadB) (Mehlhoff and Ostermeier 2023; Mehlhoff, et al. 2020). We chose these antibiotic resistance genes because their only known physiological roles are to provide resistance to their cognate antibiotics. Thus, in the absence of these antibiotics, any fitness effect of an amino acid substitution is a CFE. We constructed comprehensive site-saturation mutagenesis libraries of these genes and subjected them to a growth competition experiment (**supplementary fig. S1**, Supplementary Material online). For this experiment, we induced antibiotic gene expression in exponentially growing cultures of each library at time zero and allowed them to grow for 10 generations. We collected plasmid DNA at time zero and at 10 generations and subjected it to deep sequencing to determine the frequency of each mutation. Fitness, which corresponds to the growth rate of the bacteria, is determined from the change in frequency of the mutations relative to cells expressing the wildtype protein.

We found that the prevalence of statistically significant deleterious CFEs of mutations varied greatly among four antibiotic resistance genes: TEM-1 (21.6%), *aadB* (3.8%), *CAT-I* (0.9%), and *NDM-1* (0.2%). We chose a set of representative mutations to study in more detail. TEM-1 mutations with deleterious effects always caused TEM-1 aggregation (29/29). In addition, deleterious TEM-1 signal sequence mutations caused improper precursor processing, and deleterious mutations to and at cysteine residues caused improper inter-molecular disulfides. TEM-1’s α domain (residues 69-212) was conspicuously prone to deleterious CFEs compared to the α/β domain (residues 26-68 and 213-290). However, although mutations in the other three proteins often caused aggregation, aggregation was neither necessary nor sufficient to cause deleterious effects for those proteins (Mehlhoff and Ostermeier 2023). In summary, although mutations with deleterious CFEs often cause aggregation, the role of aggregation in CFEs is uncertain.

Our leading theory for the cause of deleterious CFEs is that the mutations create or expose misinteraction-prone motifs in the mutated protein. Exposure of motifs might come from destabilization of TEM-1’s native state as most mutations destabilize proteins (Pakula, et al. 1986). These misinteractions could be either with themselves (i.e. self-aggregation) or with other *E. coli* proteins (for example by negatively affecting the other protein’s activity through a soluble or co-aggregating interaction). As interactions increase with concentration, CFEs might offer a possible explanation for the E-R anti-correlation, complementing and overlapping with previously proposed protein-misfolding (Drummond, et al. 2005; Geiler-Samerotte, et al. 2011) and protein-misinteraction hypotheses (Yang, et al. 2012) for the E-R anticorrelation.

TEM-1’s synthesis, export, and folding pathway provide important context for understanding its potential CFE mechanisms. TEM-1 with its N-terminal signal peptide (preTEM-1) is exported to the *E. coli* periplasm in an unfolded state via the Sec pathway. In addition to directing preTEM-1 for export, the signal sequence retards TEM-1’s folding into its native state (Laminet and Pluckthun 1989). Post-translocational cleavage of preTEM-1’s signal peptide by signal peptidase I catalyzes the release of mature TEM-1 to the periplasm (van Dijl, et al. 1988). Folding of TEM-1 does not follow the basic two-state model of a simple equilibrium between unfolded and native protein. Rather, folding of TEM-1 occurs via intermediates in which the α domain is collapsed and partially folded and the two lobes of the α/β domain have not associated (Vanhove, et al. 1998). Formation of native TEM-1 from these intermediates occurs via two major parallel pathways, both involving the rate-limiting step of trans to cis isomerization of the E166-P167 bond in the Ω-loop (Vanhove, et al. 1996), in which the N- and C-terminal lobes of the α/β domain associate. Mutations that stabilize the α domain have been identified from natural isolates (Huang and Palzkill 1997; Sideraki, et al. 2001) and in lab evolution experiments (Bershtein, et al. 2008) for being beneficial to antibiotic resistance through increasing soluble TEM-1 protein expression and reducing aggregation. These mutations provide thermodynamic and/or kinetic stability, which suggests that the outcome of TEM-1 folding (native vs. aggregation) depends on the state of the α domain in these folding intermediates. This fits with our observation that the α domain is particularly prone to deleterious CFEs.

What is not clear is whether TEM-1 aggregates themselves are toxic (e.g., for the cost of their clearance, because they sequester essential *E. coli* proteins, or because they trigger stress responses with fitness costs) or whether aggregation is an unrelated consequence that shares a common cause with CFEs. One approach to establishing the role of aggregation in CFEs would be to attempt to establish a molecular mechanism for specific mutations. As a complementary approach, we instead chose to examine how TEM-1’s CFE landscape changes with 1) changes to the growth environment in ways that are expected to alter the amount of aggregation and 2) modulation of TEM-1’s stability through mutations or the addition of TEM-1 inhibitors. We hoped that patterns of how these factors impacted CFEs might provide insight into their mechanisms in part by identifying mutations that may affect fitness through different mechanisms (Schmidlin, et al. 2024).

## Results and Discussion

### Fitness effects of environmental changes that alter protein synthesis rates

Increased expression exacerbates aggregation. Accordingly, switching to a stronger promoter has been shown to increase CFEs for mutations in GFP expressed in yeast (Wu, et al. 2022). We sought to investigate aggregation’s role in TEM-1 CFEs by measuring how environmental changes that are expected to alter the amount of aggregation alter the CFE landscape. Protein scientists know that mutations often cause proteins to aggregate, which is problematic for obtaining correctly folded, active proteins to study in vitro. A common solution for reducing aggregation is to lower the growth temperature during protein production. This strategy works by reducing the protein synthesis rate, which reduces the concentration of aggregation-prone folding intermediates as well as shifts the equilibrium to the native state. If misinteractions involving TEM-1 folding intermediates (e.g., aggregation) are the primary cause of deleterious CFEs, then one would expect that lowering the growth temperature would decrease the prevalence and magnitude of deleterious CFEs. In contrast, if deleterious CFEs are primarily caused by natively folded, mutant TEM-1 misinteracting with another *E. coli* protein, the decrease in temperature would likely stabilize the weak interaction causing an increase in deleterious CFEs.

We found that reducing the growth temperature from 37°C to 30°C greatly reduced the frequency and magnitude of deleterious CFEs (**fig. 1A,C** and **2A,B**). At 30°C, only 3.7% of missense mutations caused deleterious fitness effects in both replica experiments (*P* < 0.01), compared to 21.4% at 37°C. The shift to 30°C ameliorated the deleterious effects of all classes of mutations that caused deleterious effects at 37°C (e.g., signal sequence mutations, α-domain mutations, mutations involving cysteine, and nonsense mutations).

**Figure 1.**
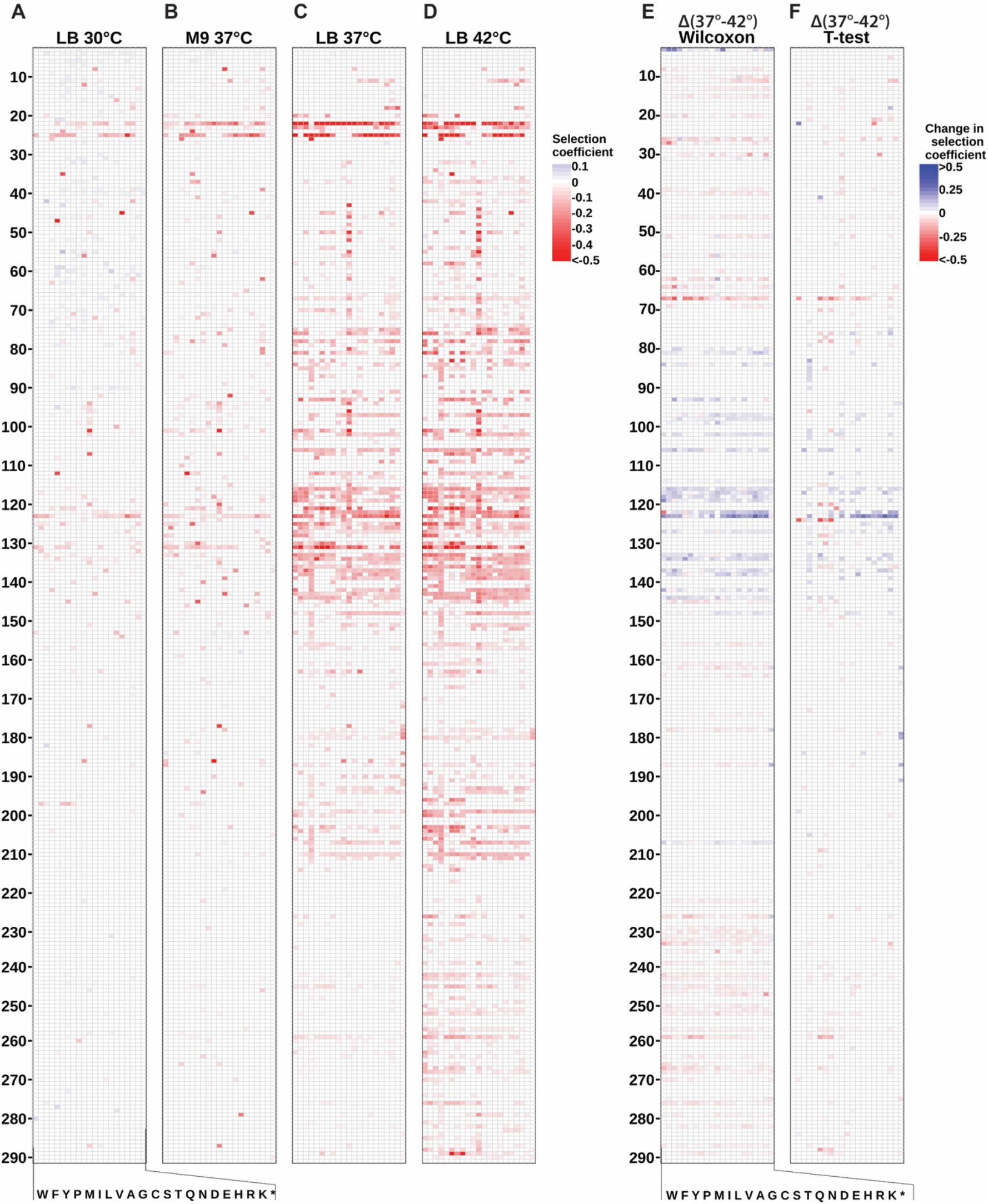
Collateral fitness effects of TEM-1 mutations under different environmental conditions. Heat map of weighted mean selection coefficients for mutations that caused fitness effects (*P* < 0.01 in both replica experiments) under four different growth conditions: **A**) LB media at 30°C, **B**) M9 minimal media at 37°C, **C**) LB media at 37°C (Mehlhoff, et al. 2020), and **D**) LB media at 42°C. Each column represents the amino acid substitution (indicated by single-letter codes at the bottom), and each row represents a position in the protein sequence (numbered on the left). Selection coefficients are relative to the wildtype under those conditions and color-coded according to the scale on the right, with blue indicating positive effects, white indicating neutral effects, and red indicating deleterious effects. Heat maps showing all selection coefficients can be found as **supplementary fig. S2-4**, Supplementary Material online. Significant changes in selection coefficient upon shift from 37°C to 42°C as evaluated by **E**) Wilcoxon signed rank analysis and **F**) Student’s t-test using data from individual codons.

Another method that protein scientists use to decrease aggregation is to switch from rich media to minimal media. Like lowering the temperature, the shift to minimal media slows the growth rate and protein synthesis rate, which reduces the concentration of aggregation-prone intermediates. However, minimal media represents a more stressful growth environment that places additional metabolic demands on cells. This increased cellular stress might exacerbate the impact of deleterious mutations, like how environmental stress can amplify the effects of underlying vulnerabilities in many biological systems (Kondrashov and Houle 1994; Stearns and Fenster 2016).

Thus, to complement our results at 30°C, we examined how changing to M9 minimal media altered CFEs. We used a modified strain that alleviated a known growth defect in M9 minimal media of DH5α-derived strains (see Materials and Methods) (Jung, et al. 2010). Like switching to 30°C, switching to M9 minimal media caused a substantial decrease in deleterious CFEs, with only 6.3% of missense mutations causing deleterious fitness effects in both replica experiments (*P* < 0.01) (**fig. 1C**). M9’s reduction in the frequency of deleterious effects was not as prominent as the reduction caused by the switch to 30°C, which is consistent with the growth rate in M9 at 37°C being faster than that in LB at 30°C (**fig. 2C**). Thus, the added stress of growing in minimal media did not exacerbate the frequency or magnitude of deleterious CFEs. Rather, the reduced growth rate ameliorated the CFEs.

**Figure 2.**
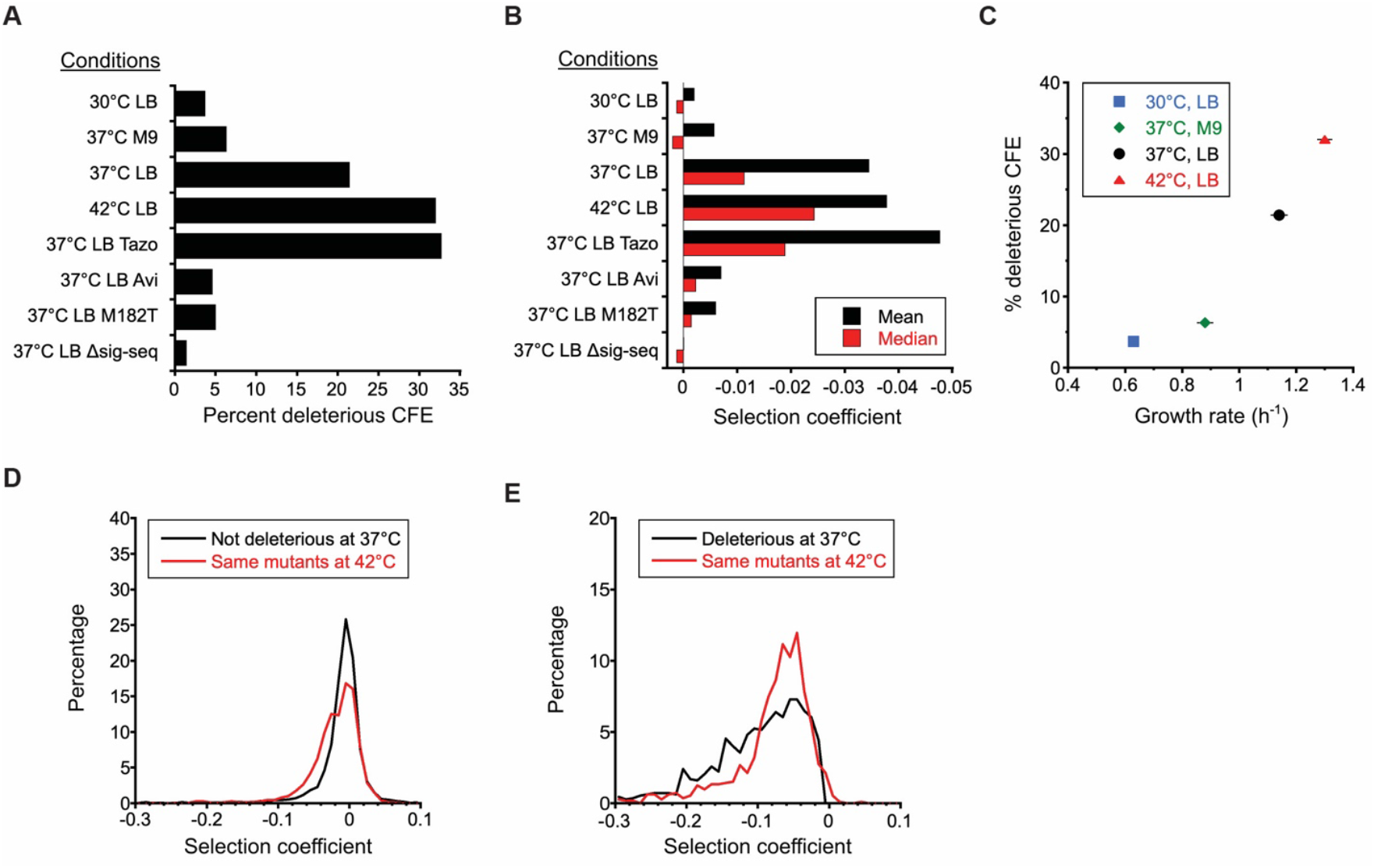
Statistics on CFE landscapes. (**A**) Frequency of deleterious CFEs (*P* < 0.01 in both replicas) and (**B**) average value of all selection coefficients under different conditions. Cells were grown in the media (LB or M9) and at the temperature 30°C, 37°C, or 42°C as indicated. Tazo, media supplemented with tazobactam; Avi, media supplemented with avibactam; M182T, TEM-1 also had the M182T mutation; Δsig-seq, TEM-1 lacked its signal sequence. Tabulated values can be found in **supplementary table S1**, Supplementary Material online. (**C**) Correlation of frequency of deleterious CFE (*P* < 0.01 in both replicas) and the growth rate of *E. coli* expressing wild-type TEM-1 under different environmental conditions. Error bars for growth rate represent standard deviation (*n* = 3-7). Change in distribution of fitness effects upon shift to 42°C for (**D**) mutations whose selection coefficient at 37°C were not significantly different than 0 (*P* > 0.01 in at least one replica) and (**E**) mutations whose selection coefficient at 37°C were significantly different than 0 (*P* < 0.01 in both replicas). The distributions at 37°C are shown in black. In red is the corresponding distribution of those same mutants at 42°C.

Raising the growth temperature from 37°C to 42°C increases the rates of growth and protein synthesis, so one might expect this shift to exacerbate deleterious CFEs. However, the shift also brings induction of the heat shock response, which includes chaperones to counteract protein misfolding. We found that shifting to 42°C caused an increase in the frequency of deleterious effects from 21.4% to 32.0% and more than a two-fold increase in the median deleterious fitness effect (**figs. 1D, 2B**). This aligns with expectations that higher temperatures would increase protein aggregation propensity and destabilize marginally stable proteins. The increase in the frequency of deleterious CFEs with temperature fits findings that the rate of evolution decelerates with temperature (Zheng, et al. 2024). Overall, across the four environmental conditions, the frequency of deleterious CFEs increases monotonically with growth rate (**fig. 2C**), consistent with the theory that TEM-1 folding intermediates cause CFEs.

However, a closer examination of how the shift to 42°C altered the fitness effect of mutations paints a more complex picture. Although switching to 42°C increased both the frequency and median deleterious effect, a significant fraction of mutations become *less* deleterious at 42°C (**fig. 1E,F** and **2D,E**). Mutations that became less deleterious clustered in the first half of the α-domain comprising helices 2-5. Mutations that became more deleterious tended to occur in the α/β-domain and the signal sequence. One interesting exception to this segregation is S124, where several mutations become more deleterious at 42°C despite being surrounded by positions that become less deleterious at 42°C (e.g., C123). Mutations to P67, which is located at the boundary between the first lobe of the α/β-domain and the α-domain, exhibited the largest increase in deleterious effect at 42°C. This temperature sensitivity suggests P67 has an important role in keeping the two subdomains of the α/β-domain associated.

A limitation in using changes in temperature and media to investigate CFEs is that these factors do more than just change the concentration of folding intermediates. Gene expression patterns and cell physiology are a function of media and temperature. Furthermore, the strength of putative misinteractions and the nature of the folding intermediates might change with temperature. Because mutational destabilization often underlies aggregation, we next examine the relationship between protein stability and CFEs.

#### CFEs correlate with protein stability for some proteins

The fact that deleterious CFEs were so rare in CAT-I and NDM-1 indicates that protein destabilization alone is insufficient to cause CFEs. However, this does not necessarily mean that protein destabilization lacks a role in deleterious CFEs. We sought to examine the relationship between protein stability and CFEs. We computationally predicted the change in the Gibbs free energy of folding (ΔΔG°_f_) of all single amino acid substitutions in our four proteins using ThermoMPNN, a state-of-the-art deep neural network trained to predict stability changes of point mutations (Dieckhaus, et al. 2024) that recently performed best on a large protein stability data set (Beltran, et al. 2025). The distributions of predicted stability effects were similar for all four proteins (**supplementary fig. S9**, Supplementary Material online). We then asked how well the stability effects predicted collateral fitness effects in TEM-1 and AadB by calculating the Spearman correlation coefficient (SCC). We chose SCC instead of Pearson’s correlation coefficient because we did not expect a linear correlation. Proteins, including TEM-1, have threshold stability (Bershtein, et al. 2006; Echave and Wilke 2017) and are likely to remain folded until deleterious effects on stability exhaust this threshold (i.e. protein folding is cooperative and sigmoidal shaped).

We first evaluated the ability of ΔΔG°_f_ to predict the PFEs of TEM-1 mutations (Firnberg, et al. 2014), as a properly folded TEM-1 is essential for enzyme activity. We found that PFEs strongly correlated with ΔΔG°_f_ with a SCC of -0.65 (**fig. 3A**). The correlation between ΔΔG°_f_ and CFEs was weaker (*ρ* = -0.38) (**fig. 3B**). However, ΔΔG°_f_ did a better job at predicting CFEs in the α-domain (*ρ* = -0.45) than in the α/β domain (*ρ* = -0.31) (**fig. 3F, I**), with similar results for CFEs at 42°C (**fig. 3G,J**). The second half of the α/β domain has very few mutations with deleterious collateral fitness effect despite being as rich with destabilizing mutations as the α-domain (**supplementary fig. S9**, Supplementary Material online). The SCC for CFEs in AadB (*ρ* = -0.20) was weak reflecting the small magnitude and lower frequency of such effects in that protein (**fig. 3D**). Across the four proteins, as CFEs become more frequent, ΔΔG°_f_’s ability to predict CFEs improved (**fig. 3K**). Overall, the results indicate that mutations predicted to destabilize TEM-1 are more likely to cause deleterious CFEs. Although destabilization is not sufficient to cause CFEs, it appears that the consequences of destabilization for CFEs differ between proteins.

**Figure 3.**
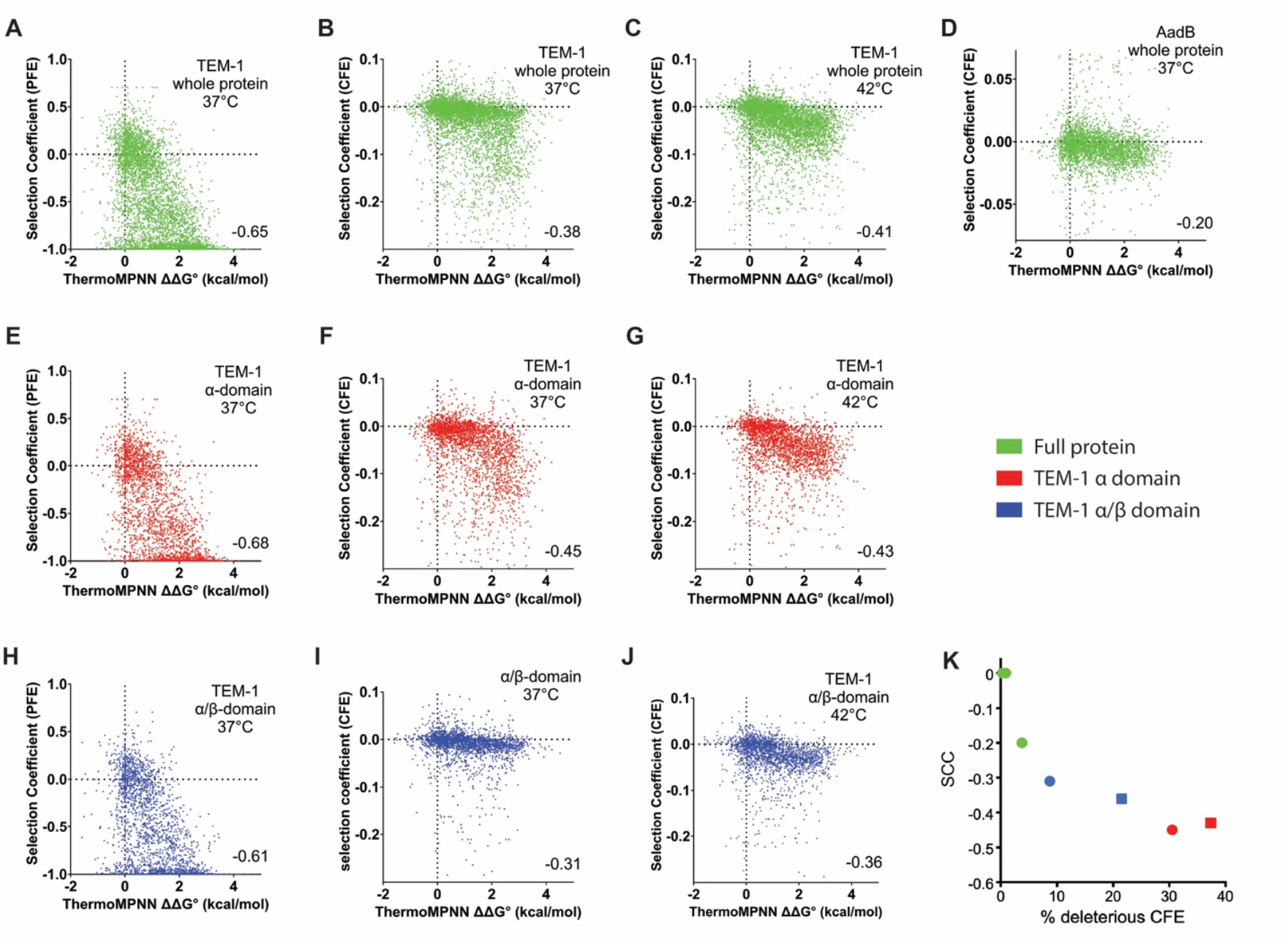
Mutational destabilization makes PFEs and CFEs more likely, especially in TEM-1’s α-domain. The correlation of CFE with ThermoMPNN predictions of ΔΔG°_f_ of mutations for **A**) PFEs of TEM-1, **B**) CFEs of TEM-1, **C**) CFEs of TEM-1 at 42°C, **D**) CFEs of AadB, **E**) PFEs of TEM-1 α-domain, **F**) CFEs of TEM-1 α-domain, **G**) CFEs of TEM-1 α-domain at 42°C, **H**) PFEs of TEM-1 α/β domain, **I**) CFEs of TEM-1 α/β domain, and **J**) CFEs of TEM-1 α/β domain at 42°C. Temperature is 37°C unless indicated otherwise. **K**) The strength of the stability-CFE correlation increases with the frequency of CFEs in the protein/domain (Green, CAT-I, NDM-1, and AadB; red, TEM-1 α-domain; blue, TEM-1 α/β domain; circles, 37°C; squares, 42°C).

#### Fitness effects of modulating protein stability

We next sought to experimentally test how TEM-1’s stability affects CFEs by two approaches: 1) using the global suppressor mutation M182T and 2) adding TEM-1 inhibitors to the growth media. Unlike the environmental changes tested above, we expected these approaches would have little to no extraneous impact on gene expression patterns and cell physiology.

The M182T mutation is a well-studied, naturally occurring TEM-1 mutation that is a global suppressor mutation that compensates for the stability loss caused by many other mutations (Huang and Palzkill 1997; Sideraki, et al. 2001). M182T stabilizes TEM-1’s native state by influencing the folding pathway (i.e. the nature of the folding intermediates) and reducing proteolysis. M182T rescues the folding defects of destabilizing mutants and confers higher thermodynamic (+1.8 kcal/mol) and kinetic stability relative to variants lacking M182T (Chattopadhyay, et al. 2022). M182T increases TEM-1’s melting temperature and soluble protein expression. While the exact mechanism of M182T’s stabilization is not yet fully elucidated, it seems to provide additional folding pathways to the native structure, particularly stabilizing TEM-1’s intermediately folded structure in which the α-domain is collapsed (Farzaneh, et al. 1996; Knies, et al. 2017; Vanhove, et al. 1998; Wang, et al. 2002) and causes a high refolding rate that allows reversible refolding (Chattopadhyay, et al. 2022).

Repeating our fitness landscape measurements in TEM-1 with M182T, we found that M182T alleviated or greatly diminished the severity of essentially all missense mutations causing deleterious CFEs (**fig. 4A,B** and **2A,B**). Only 5.0% of mutations remained deleterious in the presence of M182T (*P* < 0.01 in both replicates). Under the simplifying assumption that M182T’s 1.8 kcal/mol stabilizing effect (Chattopadhyay, et al. 2022) is additive in the context of a second mutation, M182T’s rescue of most CFEs makes sense in the context of **fig. 3B**, bringing most TEM-1 mutants’ stability into a range where CFEs are much less frequent. M182T was especially effective at rescuing even the most deleterious signal sequence mutants. M182T was less effective at rescuing the deleterious mutations to Cys in the α-domain. As these deleterious Cys mutations occur in the first half of the α-domain and caused incorrect intermolecular disulfides between two TEM-1 molecules, we have hypothesized that the incorrect disulfides form during secretion before the full α-domain is secreted and able to fold (Mehlhoff, et al. 2020). If true, it makes sense that M182T, which occurs well after these deleterious Cys mutations in the linear sequence, would not be as effective at rescuing this defect. M182T was also less effective at rescuing mutations in helices 4 and 5 in the α-domain, particularly at residues E121, L122, C123, and D131.

**Figure 4.**
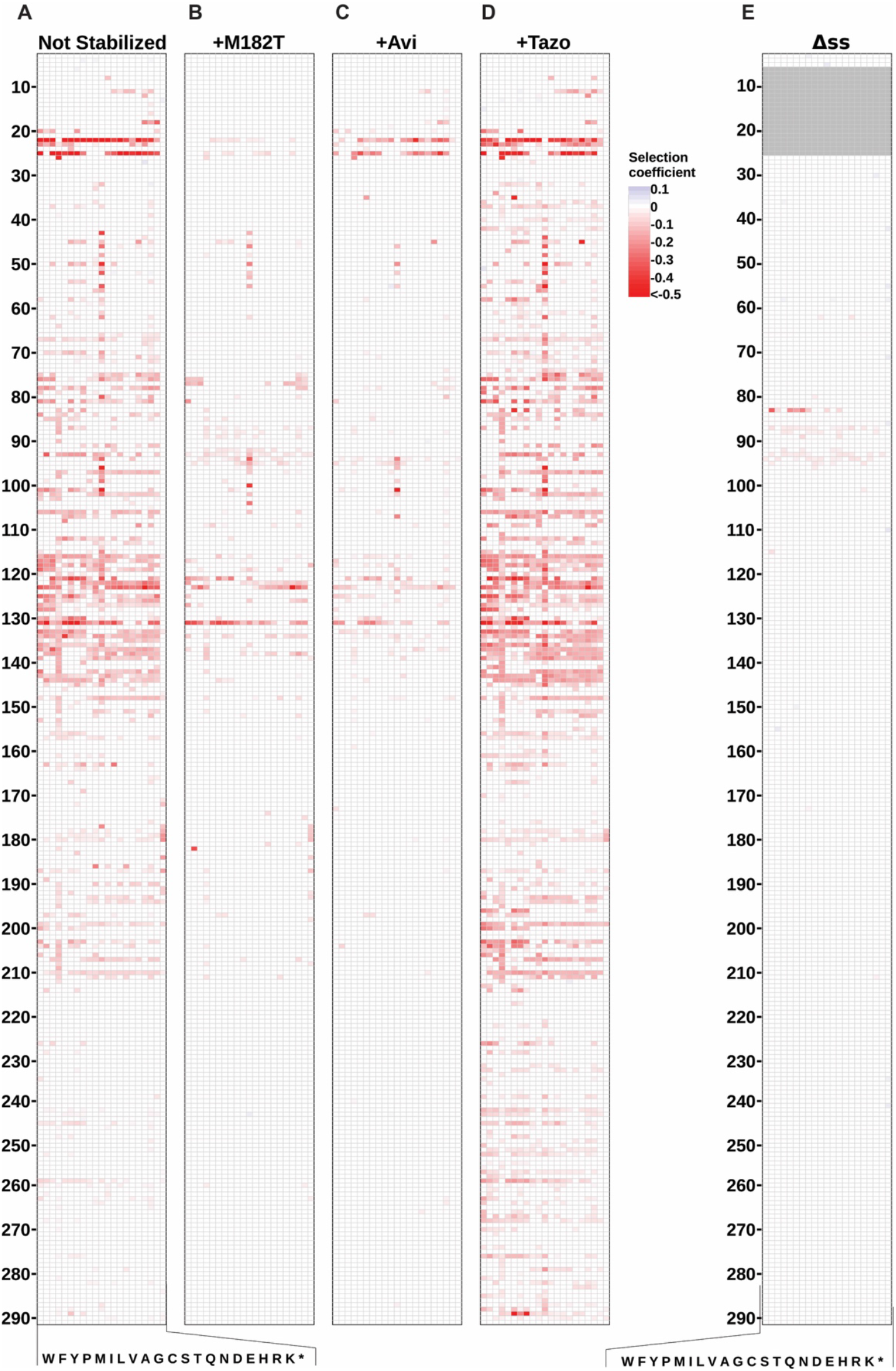
Effect of TEM-1 stability and signal sequence on collateral fitness effects of mutations. Heat map of weighted mean selection coefficients for mutations that caused fitness effects (*P* < 0.01 in both replica experiments) **A**) without a suppressor mutation or TEM-1 inhibitor (Mehlhoff, et al. 2020), **B**) with M182T suppressor mutation, **C**) with Avi, and **D**) with Tazo. All growth experiments conducted in LB media at 37°C. Each column represents the amino acid substitution (indicated by single-letter codes at the bottom), and each row represents a position in the protein sequence (numbered on the left). Selection coefficients are relative to the wildtype under those conditions and color-coded according to the scale on the right, with blue indicating positive effects, white indicating neutral effects, and red indicating deleterious effects. **E**) Heat map of weighted mean selection coefficients for mutations that caused fitness effects (*P* < 0.01 in both replica experiments) in ΔssTEM-1, a cytoplasmically-expressed variant of TEM-1 that lacks a signal sequence. Heat maps showing all selection coefficients can be found as **supplementary fig. S11-13** and **S18**, Supplementary Material online.

To complement the M182T results, we next used a non-mutational method to stabilize TEM-1. Binding to the inhibitor avibactam (Avi) stabilizes TEM-1 because binding is favorable with a *K*_d_ of 3.3 nM (Ehmann, et al. 2012) and TEM-1 binds Avi in a near-native folded state (Ji, et al. 2022). Thus, we expected the addition of Avi to the media should stabilize TEM-1 mutants (except for mutations that strongly decrease Avi affinity) and decrease the frequency and magnitude of deleterious CFEs. In contrast, tazobactam (Tazo) inhibits TEM-1 primarily via formation of a crosslinked covalent bond between catalytic residues S70 and S130 in TEM-1’s active site that disrupts TEM-1’s native structure (Maveyraud, et al. 1998; Yang, et al. 2000). Thus, we expected the addition of Tazo to the media should not rescue mutations causing deleterious CFEs and might increase the effects due to the disrupted TEM-1 structure.

We first chose a suitable saturating concentration of each inhibitor determined as the concentration that substantially reduced *E. coli’*s resistance to ampicillin but had minimal effect on *E. coli* fitness when wildtype TEM-1 was expressed in the absence of ampicillin (**supplementary fig. S10**, Supplementary Material online). We then repeated our growth competition experiment in LB at 37°C using these inhibitor concentrations (2 µg/ml Avi or 64 µg/ml Tazo) and measured the CFEs via deep sequencing as before. We found that our expectations were correct. Avi significantly decreased and Tazo significantly increased the frequency and magnitude of CFEs (**fig. 4C,D** and **2A,B**). The addition of Avi or the presence of M182T were nearly as effective at rescuing the mutations as decreasing protein synthesis rates by shifting to 30°C.

Deleterious CFEs were not rescued equally well by M182T and Avi. For example, M182T was much better than Avi at reducing the deleterious effect of signal sequence mutations (**supplementary fig. S17D**,**E**, Supplementary Material online). Also, while M182T completely rescued more deleterious mutations in the α-domain than Avi, there was a subset of mutations for which Avi was more effective, most notably many mutations at C123 and D131 as well as mutations near C77 (**supplementary fig. S17D**,**E**, Supplementary Material online). Differences in how M182T and Avi stabilize TEM-1, along with how mutations affect Avi affinity and M182T’s stabilizing effects, likely account for the rescue differences.

#### Fitness effect of changing subcellular localization

PreTEM-1 contains a signal sequence that directs the protein to be exported to the periplasm, whereupon the signal sequence is removed by signal peptidase and mature TEM-1 folds into its native structure. Deleterious TEM-1 mutations tend to activate outer envelope stress pathways (Mehlhoff, et al. 2020), suggesting that TEM-1 misinteractions in the periplasm or with the membrane cause the deleterious CFEs. Even though the reducing cytoplasm does not allow structural disulfides to form, TEM-1 lacking a signal sequence can fold into an active protein in the cytoplasm when expressed without a signal sequence (Bowden, et al. 1991) because its lone disulfide is not required for folding or catalytic activity (Laminet and Pluckthun 1989). Gross overexpression of a variant of TEM-1 in which the signal sequence is deleted results in some active cytoplasmic TEM-1 and the formation of cytoplasmic inclusion bodies. The periplasmic and cytoplasmic inclusion bodies differed in composition, sensitivity to proteases and denaturants, and morphology (the periplasmic were amorphous and the cytoplasmic were highly regular) (Bowden, et al. 1991; Valax and Georgiou 1993). Cytoplasmic TEM-1 inclusion bodies were largely free of other species, but periplasmic TEM-1 inclusion bodies included both other proteins and phospholipids (Valax and Georgiou 1993). However, these results were when TEM-1 was grossly overexpressed. Our expression level is comparable to that observed using TEM-1’s native promoter (Mehlhoff, et al. 2020).

We found that removing the signal sequence (ΔssTEM-1) was the most effective method for alleviating the deleterious CFEs of mutations (**fig. 4E** and **2A,B**). Only 1.4% of mutations caused a statistically significant fitness effect. The pattern of mutational effects was drastically different than all the periplasmic landscapes, with the most significant effects occurring at R83 and positions 87 to 95. We conclude that the signal sequence is required for most CFEs in TEM-1. Although this could indicate that the signal sequence mediates the deleterious misinteractions, we feel it more likely means that the misinteraction occurs in the periplasm and not in the cytoplasm (or if it occurs in the cytoplasm, it is not toxic there). Although we found very few deleterious mutations in ΔssTEM-1, we did observe that expression of this variant (i.e. without mutations) caused a 7.4 ± 4.8% reduction in growth rate indicating that cytoplasmic expression of ΔssTEM-1 had a fitness cost.

### Changes in deleterious collateral fitness effects correlate poorly with the extent of TEM-1 aggregation

We selected several mutations to study in detail. We confirmed the fitness effect of these mutations by measuring the growth rate of cells expressing the mutants compared to cells expressing unmutated TEM-1. These included mutants that were most deleterious at 42°C (**supplementary fig. S19A**, Supplementary Material online), mutants that were most deleterious at 37°C (**supplementary fig. S19B**, Supplementary Material online), mutants that were rescued by Avi and M182T (**supplementary fig. S20A and S21**, Supplementary Material online), mutants that were not rescued by Avi and M182T (**supplementary fig. S20B**, Supplementary Material online), and mutants whose selection coefficients decreased or increased upon removal of the signal sequence (**supplementary fig. S22**, Supplementary Material online).

We next lysed the cells and fractionated the proteins into soluble and insoluble fractions. These fractions were separated by SDS-PAGE and analyzed by western blot using anti-TEM-1 antibodies to learn the effect of the environmental/mutational changes on the amount of soluble and insoluble TEM-1. For wildtype TEM-1, the temperature shift did not appreciably alter the levels of soluble TEM-1 and we observed very little, if any, insoluble TEM-1 at any temperature (**fig. 5E, supplementary fig. S24**, Supplementary Material online). However, a faint amount of preTEM-1 in the insoluble fraction appeared at 42°C, perhaps suggesting that the rate of signal sequence cleavage was beginning to reach the point at which it could not keep up with the higher rate of protein synthesis.

**Figure 5.**
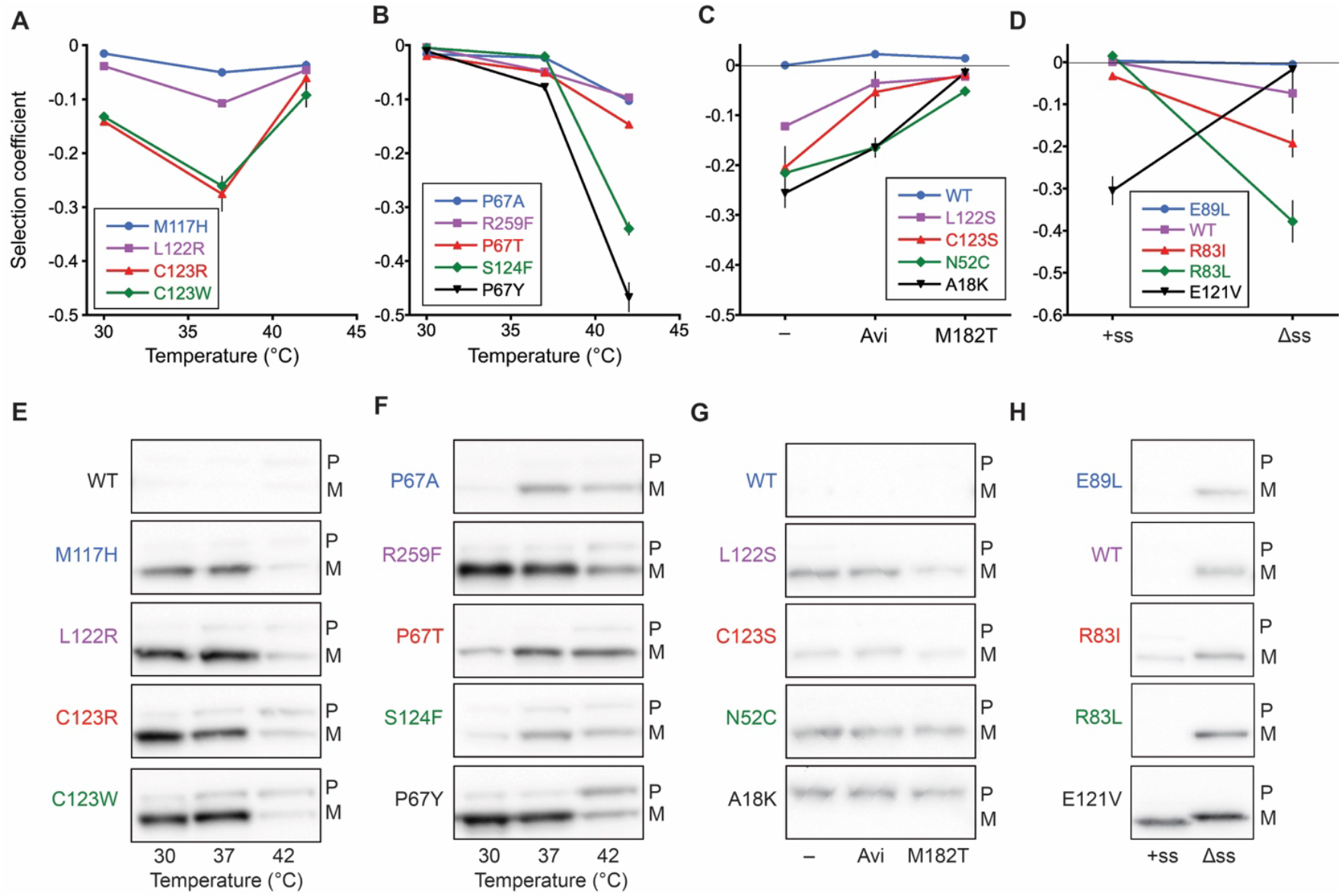
Selection coefficients and western blot analysis for select mutants. Selection coefficient for cells expressing mutants that **A**) are most deleterious at 37°C, **B**) are most deleterious at 42°C, **C**) are rescued by stabilization of TEM-1 by Avi or M182T (– = no stabilizer), **D**) have different fitness effects in the presence (+ss) and absence (Δss) of a signal sequence. Selection coefficients were measured by monoculture growth experiments. Lines are a guide for the eye. Error bars are the standard deviation (*n* ≥ 3). Western blots with TEM-1 antisera of the insoluble fractions of cells expressing mutants that **E**) are most deleterious at 37°C, **F**) are most deleterious at 42°C, **G**) are rescued by stabilization of TEM-1 by Avi or M182T (– = no stabilizer), and **H**) have different effects in the presence (+ss) and absence (Δss) of a signal sequence. Selection coefficients and full western blots of both replica experiments including the soluble fraction for these and additional mutants can be found in **supplementary figs. S19-26** and **Tables S2-4**, Supplementary Material online.

For all mutants studied for the effect of temperature, the level of soluble protein increased or remained the same as the temperature dropped from 37°C to 30°C and decreased with the shift from 37°C to 42°C (**supplementary fig. S23-24**, Supplementary Material online). In contrast, the temperature-dependent patterns of insoluble mutant TEM-1 varied greatly. For some mutants, the patterns were consistent with the hypotheses that magnitude of the deleterious CFEs increases with the amount of insoluble TEM-1 and that temperature alters the magnitude of the deleterious effects by altering the level of insoluble protein. However, in many cases, they were not. For example, among the mutations that were most deleterious at 37°C, all showed less insoluble TEM-1 at 42°C; however, for many of these mutants, the level of insoluble TEM-1 at 30°C was comparable to that at 37°C even though the deleterious effects were substantially diminished at 30°C (**fig. 5A,E**). Similarly, R259F and R259I became increasingly deleterious with temperature, but the level of insoluble TEM-1 did not increase. Furthermore, even mutations at the same position had different patterns (**fig. 5B,F**). S124V and S124F become markedly (and equally) deleterious at 42°C; however, S124V exhibited an increase of insoluble TEM-1 at 42°C and S124F did not (**supplementary fig. S23-24**, Supplementary Material online). Among the mutations at P67 that become much more deleterious at 42°C, P67A and P67T showed little change in insoluble TEM-1 but P67Y, the most deleterious of the P67 mutations, showed a decrease in insoluble TEM-1 but a distinct increase in insoluble preTEM-1.

A similar disconnect exists between how M182T and especially Avi modulated the deleterious fitness effects and the extent of aggregation. Mutants that were robustly rescued by M182T (A18K, N52C, L122S, and C123S) showed marginal to no reduction in insoluble TEM-1 or preTEM-1 (**fig. 5C,G**). Avi failed to reduce the amount of insoluble TEM-1 for any mutant for which it reduced the severity of the CFEs. Mutants that were not rescued by M182T (G78R and E121V) also showed little change in insoluble protein (**supplementary fig. S25B**, Supplementary Material online). M182T and Avi exacerbated the inhibition of signal sequence cleavage caused by A18K, despite both rescuing A18K’s fitness defect (**supplementary fig. S25A**, Supplementary Material online).

We also found a lack of a clear trend between CFEs and the amount of insoluble TEM-1 with removal of the signal sequence (**fig. 5D,H, supplementary fig. S26**, Supplementary Material online). Removal of TEM-1’s signal sequenced from WT, R83I, and R83L caused reductions in fitness and caused TEM-1 to aggregate (soluble TEM-1 was also present), though it was not apparent that R83I and R83L caused more aggregation than WT despite them being significantly more deleterious. Removal of the signal sequence from E121V did not alter the amount of insoluble TEM-1 despite rescuing the fitness defect caused by expressing the E121V protein in the periplasm.

#### Perspective

A recent preprint by Quan et al. also predicted stability effects of mutations in our four proteins (and three additional proteins) and concluded that CFEs do not correlate with predicted destabilization (Quan, et al. 2025), in contrast to our findings here for TEM-1. Differences in approach and interpretation likely explain the different conclusions. Quan et al. primarily predicted stability effects using FoldX, whereas we used ThermoPNN, a more recent program that performed better against benchmarks in direct comparisons (Beltran, et al. 2025; Dieckhaus, et al. 2024). Quan et al. evaluated a linear correlation using PCC, whereas we used SCC because proteins in general and TEM-1 in particular exhibit threshold stability (Bershtein, et al. 2006; Echave and Wilke 2017). We agree that destabilization does not always lead to deleterious CFEs and that CFEs need not require destabilization, but our computations and experiments illustrate that 1) destabilizing TEM-1 mutations are more likely to cause deleterious CFEs and 2) most deleterious TEM-1 mutations can be rescued by mutations or ligands that stabilize TEM-1. Both studies highlight that the consequences of destabilization for CFEs is protein-dependent, and that the extent of aggregation does not correlate well with the magnitude of deleterious CFEs, even for a protein like TEM-1 for which CFEs and stability correlate.

Both reducing the growth rate and stabilizing TEM-1 ameliorated most deleterious CFEs. Both, in general, would be expected to decrease aggregation. However, increasing the growth rate by increasing the growth temperature increased the deleterious effects for some mutants but ameliorated it for others. Furthermore, the extent of aggregation correlated with the magnitude of CFEs for only a few of the mutations/conditions tested. Overall, the complexity of the patterns of fitness effects and the amount of aggregation defies the simple explanation that more insoluble protein typically leads to larger deleterious CFEs.

We offer a few speculative views. First, TEM-1 with deleterious mutations must interact with some “target” native *E. coli* molecule to affect fitness. This might be a direct harm, such as compromising a cellular process with fitness implications, or an indirect harm, such as inducing a stress response with fitness costs or costing the cell resources to remove aggregates. Second, the mutation must create, strengthen, or increase the exposure of a protein motif that interacts with this target molecule (or another TEM-1 molecule). Third, destabilizing mutations are the easiest route to increasing exposure of existing or mutant TEM-1 motifs to the cellular contents, especially if they are inaccessible when TEM-1 is natively folded and stable. Destabilization also commonly leads to aggregation, but aggregation motifs and CFEs motifs need not be the same motif; destabilization may be the common cause for different phenomena. Fourth, although the overall extent of aggregation often fails to predict the magnitude of the deleterious effect, we should not expect that such aggregates are comprised of a single species aggregating in the same manner. Some species may be detrimental, and some may be benign. The frequency of detrimental species may be mutation- and stability-dependent. Fifth, the activation of periplasmic stress responses by deleterious mutations (and lack of induction cytoplasmic stress responses) (Mehlhoff, et al. 2020) coupled with the nearly complete rescue of deleterious effects by changing expression from the periplasm to the cytoplasm strongly suggests the target molecule(s) that TEM-1 interacts with is in the periplasm or membrane.

## Materials & Methods

### Strains, plasmids, chemicals, and growth conditions

For all experiments (except those in M9 minimal media) the strain was NEB 5-alpha LacIq (F′ proA+B+ lacIq Δ(lacZ)M15 zzf::Tn10 (TetR) / fhuA2Δ(argF-lacZ)U169 phoA glnV44 Φ80Δ(lacZ)M15 gyrA96 recA1 relA1 endA1 thi-1 hsdR17). For M9 minimal media we used a variant we isolated of NEB 5-alpha LacIq in which its mutant *purB* gene (E115K) had reverted to wildtype *purB* (K115E). The E115K *purB* mutation confers a higher transformation efficiency but also a slower growth in minimal media as the gene catalyzes two reactions in the de novo biosynthesis pathway for AMP formation (Jung, et al. 2010). We found that the *purB* (E115K) gene in NEB 5-alpha LacIq was prone to spontaneous K115E or K115Q mutations that increased growth rate in M9 minimal media, which complicated the growth competition experiments with the library. Performing the experiment with our isolated NEB 5-alpha LacIq purB(K115E) strain solved this problem.

The TEM-1 gene was under the control of the IPTG-inducible tac promoter on pSKunk1, a minor variant of plasmid pSKunk3 (AddGene plasmid #61531) (Firnberg and Ostermeier 2012). Initial MIC experiments with avibactam were performed with avibactam from VWR/Avantor (75837-390). All other experiments were with avibactam from MedChemExpress (HY-14879A). Tazobactam was from Selleck Chemicals (S3077) and purchased from VWR/Avantor. All growth experiments were conducted in LB media or M9 minimal media. All media was supplemented with glucose (2% w/v) and spectinomycin (50 µg/ml) to maintain the pSKunk1 plasmid except where otherwise noted. Expression of TEM-1 was induced by addition of 1 mM IPTG. Experiments used 100 ml media in 500 ml baffled flasks with shaking at 250 rpm. The optical density (OD) of cultures was measured at 600 nm.

### TEM-1 libraries

We used our previously constructed library of all the possible single-codon substitutions (i.e. 5’-NNN) in TEM-1 (Mehlhoff, et al. 2020). This library was constructed in three regions due to the read length of the deep sequencing. In this work, libraries containing the M182T mutation or in TEM-1 lacking the signal sequence (ΔssTEM-1) were created by the same method. The ΔssTEM-1 construct was the same as described previously (Bowden, et al. 1991). The 5’ region of the coding region was 5’-ATG.CGT.TTT.CAC. This encodes Met.Arg.Phe.His with the His being the first amino acid of the native, mature TEM-1.

In replica 1 of the TEM-1(M182T) library, we noticed underrepresented mutations at a few positions within regions 1 and 3 of the library. We created a supplemental library containing NNN mutations at V31, L191, L196, G226, W227, F228, S233, I244, and G251. This supplemental library was spiked into the original library in the replica 2 growth competition experiment.

### Growth competition for measurements of fitness effects

Fitness was measured by deep mutational scanning as described (Mehlhoff and Ostermeier 2023) using a growth competition experiment. Briefly, expression of TEM-1 in the exponentially growing library cultures was induced with IPTG, and the culture was allowed to grow for approximately ten generations with a single dilution at about five generations. Plasmid DNA was collected from before induction and after 10 generations of growth. Custom adapters were added to the plasmid DNA by PCR. Adapters were designed to be compatible with the Illumina platform and contained barcodes for identification of each timepoint and sample. Proper DNA size of the PCR products was validated by agarose electrophoresis gel before the PCR products were pooled and submitted for Illumina MiSeq (2 x 300 bp reads) at the Single Cell and Transcriptomics Core facility at Johns Hopkins University.

### Deep sequencing analysis

Illumina MiSeq reads for each of the three regions were inspected for per base sequence quality using FastQC (Wingett and Andrews 2018). Paired-end reads were merged using PEAR (Zhang, et al. 2014) set to a minimum assembly length of 200 base pairs. Illumina adapters and base pairs outside the desired regions were cropped from the merged reads using Trimmomatic (Bolger, et al. 2014) (Region 1-HEADCROP:24, CROP:285; Region 2-HEADCROP:20, CROP:285; Region 3-HEADCROP:24, CROP:291). The resulting trimmed reads were input to Enrich2 (Rubin, et al. 2017), which counted the variants for use in calculating selection coefficients and variance. Reads containing bases with a quality score below 20, bases marked as N, or mutations at more than one codon were filtered out.

#### Fitness calculation

Fitness of an allele (*w*_*i*_) was calculated from the enrichment of the synonyms of the wild-type gene (*ε*_*wt*_), the enrichment of allele *i* (*ε*_*i*_) and the fold increase in the number of cells during the growth competition experiment (*r*) as described by Equation 1.

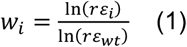

We utilize the frequency of synonyms of the wild-type gene as the reference instead of the frequency of wildtype because wildtype synonyms occurred more frequently in the library and wildtype sequencing counts are more prone to being affected by the artifact of PCR template jumping during the preparation of barcoded amplicons for deep sequencing. We defined the enrichment of allele *i* as

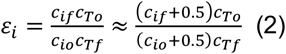

which was calculated from the counts of that allele (*c*_*i*_) and the total sequencing counts (*c*_*T*_) in which the subscripts *o* and *f* refer to the beginning and end of the experiment, respectively. We add the 0.5 to all counts of mutant alleles to assist with alleles that have counts that are zero (i.e. so that a fitness value can still be calculated). We calculated the variance in the fitness (w) from Equation 3’

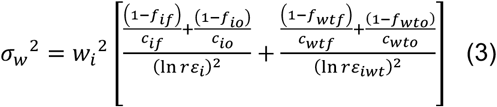

where the frequency of allele (*f*_*i*_) was calculated from Equation 4.

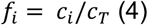

From the variance in fitness, we calculated a 99% confidence interval. Additionally, we calculated a *p*-value using a 2-tailed test. Detailed derivations and explanations of all calculations can be found in our previous work (Mehlhoff and Ostermeier 2020; Mehlhoff, et al. 2020). Selection coefficients were defined as *w* -1.

### Wilcoxon signed rank test analysis

We employed Wilcoxon signed rank tests to evaluate if the changes in fitness of mutations at a position tended to be distributed to be more deleterious or less deleterious than expected at random upon the change in condition than expected at random (e.g., temperature, presence of Avi/M182T). On the Wilcoxon landscape heatmaps, we color all mutations with the change in weighted mean selection coefficients at that position if the trend is significant (*p*<0.01)

### Student’s t-test using individual codons

We used a Student’s t-test to analyze whether an amino acid substitution has a different fitness effect between two different conditions. From our two replica experiments, we have multiple independent measures of the fitness effect of an amino acid substitution – one for each codon of that amino acid. For the t-test, we consider each fitness measurement for a codon to be a different measure of the effect of that amino acid substitution. For example, for a glycine substitution there are 4 codons x 2 replica experiments giving 8 measures of the effect of a glycine substitution. We recognize that this is not the same as 8 biological replicas, but it is something more than only two replicas. It also means the power to measure significant effects depends on the number of codons for that amino acid. Nonetheless, we found this method useful to identify mutations with likely condition-dependent differences in fitness effects.

### ThermoMPNN calculations

We performed ΔΔG°_f_ predictions by ThermoMPNN (Dieckhaus, et al. 2024) using a copy of the Google Colaboratory notebook provided by the Kuhlman Lab https://github.com/Kuhlman-Lab/ThermoMPNN-D/blob/ThermoMPNN-I/ThermoMPNN-I.ipynb. The structures used for the calculations were 1AXB for TEM-1 (Maveyraud, et al. 1998), 4WQK for AadB (Cox, et al. 2015), 3U9B for CAT-I (Biswas, et al. 2012), and 7UOX for NDM-1 (Mandal, et al. 2022).

### Minimum Inhibitory Concentration (MIC) Assays

Cultures were grown overnight at 37°C in LB media containing 2% w/v glucose and 50 μg/mL spectinomycin. The overnight cultures were then diluted 10-fold to an OD_600_ between 0.050 and 0.500. The concentration of cells was calculated by multiplying the OD_600_ by 8 × 10^8^. A total of 3 × 10^6^ cells were added to 3 mL LB media containing 2% w/v glucose, 50 μg/mL spectinomycin, and 1 mM IPTG. Differing levels of antibiotic and inhibitor were added to each tube. Cultures were incubated for 14-18 h at 37°C in a shaker set to 250 rpm before the OD_600_ were measured in triplicate. Cultures with an OD600 greater than 0.500 were diluted 10-fold and re-measured. We identified the MIC as the lowest inhibitor concentration at which the OD_600_ of cultures with 25 μg/mL ampicillin were not statistically different from the OD_600_ of media not inoculated with cells by a Student’s t-test (*P* < 0.05).

### Monoculture growth assay for fitness

Cultures were grown following the same protocol described by Mehlhoff *et al*. (Mehlhoff, et al. 2020). Briefly, we tracked the optical density of monocultures over six hours of induced growth with a dilution after about three hours. We calculated the fitness using Equation 5 from the dilution factor (*d*) and the starting and final ODs (*O*_*o*_, *O*_*f*_) of the mutant and wild-type cultures under the assumption that the correlation between OD and cell density was the same for cells whether they were expressing wild-type or any mutant TEM-1 allele. Thus, fitness represents the mean growth rate of the cells expressing the mutant protein relative to the growth rate of cells expressing the wildtype protein.

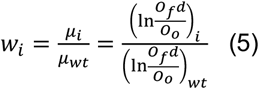

### Cell fractionation and analysis by PAGE and western blot

Cell fractionation and, SDS-PAGE, and Western blots were performed essentially as previously described (Mehlhoff, et al. 2020). Briefly, cultures were grown as in the monoculture growth assays. Ten generations post induction of TEM-1 expression, cultures were lysed using BugBuster and fractionated into soluble and insoluble fractions by centrifugation. After SDS-PAGE electrophoresis of these samples, TEM-1 was detected by western blot using anti-β-lactamase antibodies.

## Acknowledgements

This research was supported by National Science Foundation grants MCB-1817646 and MCB-2113019 to M.O. Authors have no competing interests.

## References

Beltran A, Jiang X, Shen Y, Lehner B 2025. Site-saturation mutagenesis of 500 human protein domains. Nature 637: 885–894.

Bershtein S, Goldin K, Tawfik DS 2008. Intense neutral drifts yield robust and evolvable consensus proteins. J Mol Biol 379: 1029–1044.

Bershtein S, Segal M, Bekerman R, Tokuriki N, Tawfik DS 2006. Robustness-epistasis link shapes the fitness landscape of a randomly drifting protein. Nature 444: 929–932.

Biswas T, Houghton JL, Garneau-Tsodikova S, Tsodikov OV 2012. The structural basis for substrate versatility of chloramphenicol acetyltransferase CATI. Protein Sci 21: 520–530.

Bolger AM, Lohse M, Usadel B 2014. Trimmomatic: a flexible trimmer for Illumina sequence data. Bioinformatics 30: 2114–2120.

Bowden GA, Paredes AM, Georgiou G 1991. Structure and morphology of protein inclusion bodies in Escherichia coli. Biotechnology (N Y) 9: 725–730.

Chattopadhyay G, Bhowmick J, Manjunath K, Ahmed S, Goyal P, Varadarajan R 2022. Mechanistic insights into global suppressors of protein folding defects. PLoS Genet 18: e1010334.

Cox G, Stogios PJ, Savchenko A, Wright GD 2015. Structural and molecular basis for resistance to aminoglycoside antibiotics by the adenylyltransferase ANT(2’’)-Ia. MBio 6.

Dieckhaus H, Brocidiacono M, Randolph NZ, Kuhlman B 2024. Transfer learning to leverage larger datasets for improved prediction of protein stability changes. Proc Natl Acad Sci U S A 121: e2314853121.

Drummond DA, Bloom JD, Adami C, Wilke CO, Arnold FH 2005. Why highly expressed proteins evolve slowly. Proc Natl Acad Sci U S A 102: 14338–14343.

Echave J, Wilke CO 2017. Biophysical Models of Protein Evolution: Understanding the Patterns of Evolutionary Sequence Divergence. Annu Rev Biophys 46: 85–103.

Ehmann DE, Jahic H, Ross PL, Gu RF, Hu J, Kern G, Walkup GK, Fisher SL 2012. Avibactam is a covalent, reversible, non-beta-lactam beta-lactamase inhibitor. Proc Natl Acad Sci U S A 109: 11663–11668.

Farzaneh S, Chaibi EB, Peduzzi J, Barthelemy M, Labia R, Blazquez J, Baquero F 1996. Implication of Ile-69 and Thr-182 residues in kinetic characteristics of IRT-3 (TEM-32) beta-lactamase. Antimicrob Agents Chemother 40: 2434–2436.

Firnberg E, Labonte JW, Gray JJ, Ostermeier M 2014. A comprehensive, high-resolution map of a gene’s fitness landscape. Mol Biol Evol 31: 1581–1592.

Firnberg E, Ostermeier M 2012. PFunkel: efficient, expansive, user-defined mutagenesis. PLoS ONE 7: e52031.

Geiler-Samerotte KA, Dion MF, Budnik BA, Wang SM, Hartl DL, Drummond DA 2011. Misfolded proteins impose a dosage-dependent fitness cost and trigger a cytosolic unfolded protein response in yeast. Proc Natl Acad Sci U S A 108: 680–685.

Huang W, Palzkill T 1997. A natural polymorphism in beta-lactamase is a global suppressor. Proc Natl Acad Sci U S A 94: 8801–8806.

Ji Z, Kozuch J, Mathews, II, Diercks CS, Shamsudin Y, Schulz MA, Boxer SG 2022. Protein electric fields enable faster and longer-lasting covalent inhibition of beta-lactamases. J Am Chem Soc 144: 20947–20954.

Jung SC, Smith CL, Lee KS, Hong ME, Kweon DH, Stephanopoulos G, Jin YS 2010. Restoration of growth phenotypes of Escherichia coli DH5alpha in minimal media through reversal of a point mutation in purB. Appl Environ Microbiol 76: 6307–6309.

Knies J, Cai F, Weinreich DM 2017. Enzyme efficiency but not thermostability drives cefotaxime resistance evolution in TEM-1 beta-lactamase. Mol Biol Evol 34: 1040–1054.

Kondrashov AS, Houle D 1994. Genotype-environment interactions and the estimation of the genomic mutation rate in Drosophila melanogaster. Proc Biol Sci 258: 221–227.

Laminet AA, Pluckthun A 1989. The precursor of beta-lactamase: purification, properties and folding kinetics. Embo J 8: 1469–1477.

Mandal M, Xiao L, Pan W, Scapin G, Li G, Tang H, Yang SW, Pan J, Root Y, de Jesus RK, Yang C, Prosise W, Dayananth P, Mirza A, Therien AG, Young K, Flattery A, Garlisi C, Zhang R, Chu D, Sheth P, Chu I, Wu J, Markgraf C, Kim HY, Painter R, Mayhood TW, DiNunzio E, Wyss DF, Buevich AV, Fischmann T, Pasternak A, Dong S, Hicks JD, Villafania A, Liang L, Murgolo N, Black T, Hagmann WK, Tata J, Parmee ER, Weber AE, Su J, Tang H 2022. Rapid Evolution of a Fragment-like Molecule to Pan-Metallo-Beta-Lactamase Inhibitors: Initial Leads toward Clinical Candidates. J Med Chem 65: 16234–16251.

Maveyraud L, Pratt RF, Samama JP 1998. Crystal structure of an acylation transition-state analog of the TEM-1 beta-lactamase. Mechanistic implications for class A betalactamases. Biochemistry 37: 2622–2628.

Mehlhoff JD, Ostermeier M 2020. Biological fitness landscapes by deep mutational scanning. Methods Enzymol 643: 203–224.

Mehlhoff JD, Ostermeier M 2023. Genes vary greatly in their propensity for collateral fitness effects of mutations. Mol Biol Evol 40: msad038.

Mehlhoff JD, Stearns FW, Rohm D, Wang B, Tsou EY, Dutta N, Hsiao MH, Gonzalez CE, Rubin AF, Ostermeier M 2020. Collateral fitness effects of mutations. Proc Natl Acad Sci U S A 117: 11597–11607.

Pakula AA, Young VB, Sauer RT 1986. Bacteriophage lambda cro mutations: effects on activity and intracellular degradation. Proc Natl Acad Sci U S A 83: 8829–8833.

Pal C, Papp B, Hurst LD 2001. Highly expressed genes in yeast evolve slowly. Genetics 158: 927–931.

Quan N, Eguchi Y, Brown A, Geiler-Samerotte K 2025. Collateral fitness effects of mutation are not commonly caused by protein misfolding. bioRxiv: 2025.2009.2012.675869.

Rubin AF, Gelman H, Lucas N, Bajjalieh SM, Papenfuss AT, Speed TP, Fowler DM 2017. A statistical framework for analyzing deep mutational scanning data. Genome Biol 18: 150.

Schmidlin K, Apodaca S, Newell D, Sastokas A, Kinsler G, Geiler-Samerotte K 2024. Distinguishing mutants that resist drugs via different mechanisms by examining fitness tradeoffs. Elife 13.

Sideraki V, Huang W, Palzkill T, Gilbert HF 2001. A secondary drug resistance mutation of TEM-1 beta-lactamase that suppresses misfolding and aggregation. Proc Natl Acad Sci U S A 98: 283–288.

Stearns FW, Fenster CB 2016. Fisher’s geometric model predicts the effects of random mutations when tested in the wild. Evolution 70: 495–501.

Valax P, Georgiou G 1993. Molecular characterization of beta-lactamase inclusion bodies produced in Escherichia coli. 1. Composition. Biotechnol Prog 9: 539–547.

van Dijl JM, Smith H, Bron S, Venema G 1988. Synthesis and processing of Escherichia coli TEM-beta-lactamase and Bacillus licheniformis alpha-amylase in E. coli: the role of signal peptidase I. Mol Gen Genet 214: 55–61.

Vanhove M, Lejeune A, Guillaume G, Virden R, Pain RH, Schmid FX, Frere JM 1998. A collapsed intermediate with nonnative packing of hydrophobic residues in the folding of TEM-1 beta-lactamase. Biochemistry 37: 1941–1950.

Vanhove M, Raquet X, Palzkill T, Pain RH, Frere JM 1996. The rate-limiting step in the folding of the cis-Pro167Thr mutant of TEM-1 beta-lactamase is the trans to cis isomerization of a non-proline peptide bond. Proteins 25: 104–111.

Wang X, Minasov G, Shoichet BK 2002. Evolution of an antibiotic resistance enzyme constrained by stability and activity trade-offs. J Mol Biol 320: 85–95.

Wingett SW, Andrews S 2018. FastQ Screen: A tool for multi-genome mapping and quality control. F1000Res 7: 1338.

Wu Z, Cai X, Zhang X, Liu Y, Tian GB, Yang JR, Chen X 2022. Expression level is a major modifier of the fitness landscape of a protein coding gene. Nat Ecol Evol 6: 103–115.

Yang JR, Liao BY, Zhuang SM, Zhang J 2012. Protein misinteraction avoidance causes highly expressed proteins to evolve slowly. Proc Natl Acad Sci U S A 109: E831–840.

Yang Y, Janota K, Tabei K, Huang N, Siegel MM, Lin YI, Rasmussen BA, Shlaes DM 2000. Mechanism of inhibition of the class A beta -lactamases PC1 and TEM-1 by tazobactam. Observation of reaction products by electrospray ionization mass spectrometry. J Biol Chem 275: 26674–26682.

Zhang J, Kobert K, Flouri T, Stamatakis A 2014. PEAR: a fast and accurate Illumina Paired-End reAd mergeR. Bioinformatics 30: 614–620.

Zhang J, Yang JR 2015. Determinants of the rate of protein sequence evolution. Nat Rev Genet 16: 409–420.

Zheng J, Guo N, Huang Y, Guo X, Wagner A 2024. High temperature delays and low temperature accelerates evolution of a new protein phenotype. Nat Commun 15: 2495.

