## Supplemental Material for "Environmental and mutational modulation of collateral fitness effects informs their mechanisms"

**This PDF file includes:**

Supplementary Figs S1-26
Supplementary Tables S1-4

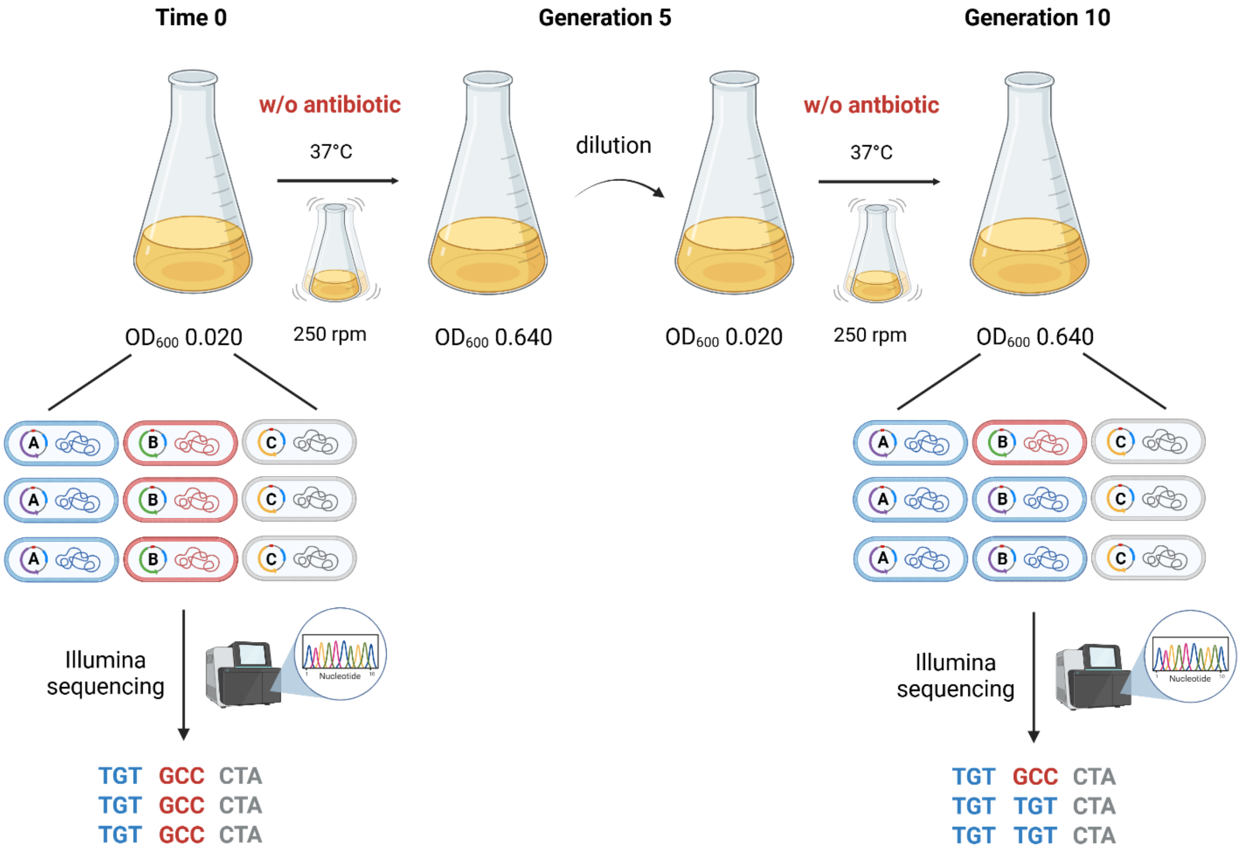


**Figure S1.** Schematic of the growth competition experiments with deep mutational scanning. TEM-1 expression was induced with IPTG at time 0. Cultures were grown for five generations from an OD of 0.020 to 0.640 before being diluted and allowed to grow for an additional five generations. Cells were pelleted and plasmid prepped for deep sequencing immediately before induction and after 10 generations of induced growth. Figure adapted from Mehlhoff et al. (Mehlhoff and Ostermeier 2020) using BioRender.com.

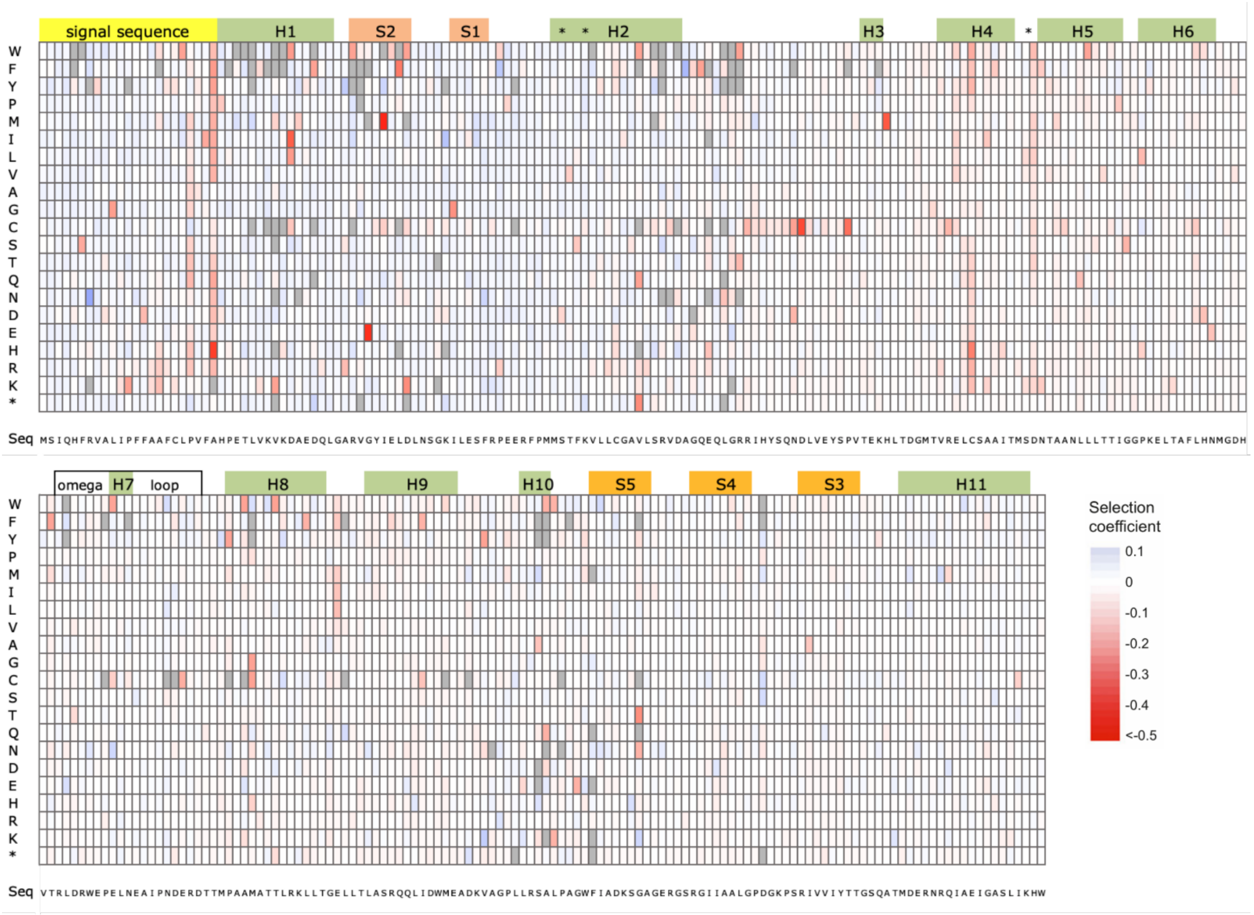


**Figure S2. Collateral fitness effects of mutations in TEM-1 at 30°C in LB media**. Heat map shows the weighted mean selection coefficient from two biological replicates. All selection coefficient values are shown regardless of whether they are significantly different than 0. Mutations with no fitness measurement in either replicate are shown in grey. Regions corresponding to the signal sequence (yellow), α-helices (green), β-strands (orange), the Ω-loop (white) and key active site residues (*) are shown above the heat map. Positions with a statistically significant difference in selection coefficient compared to 37°C in LB media can be found in **Fig S5**.

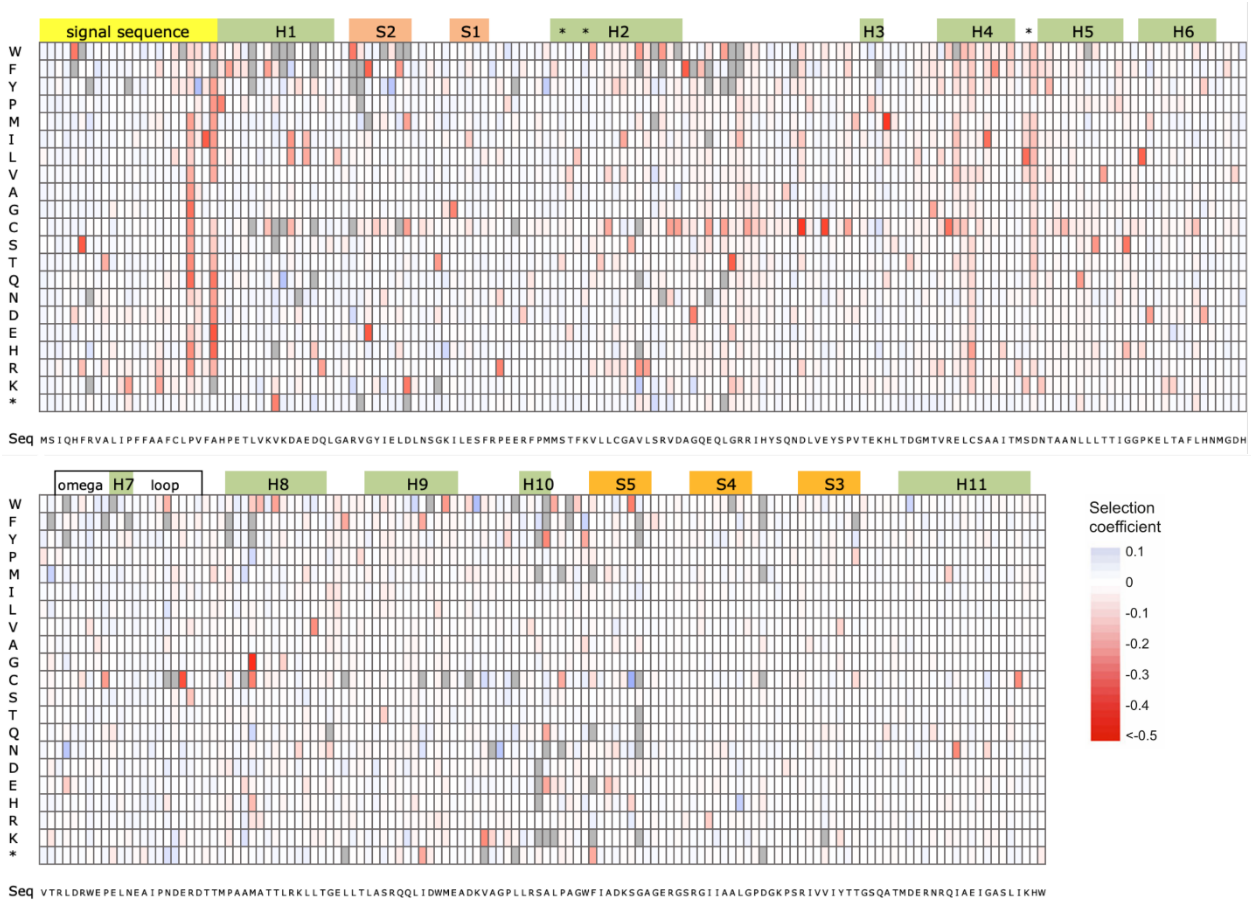


**Figure S3. Collateral fitness effects of mutations in TEM-1 in M9 minimal media at 37°C**. Heat map shows the weighted mean selection coefficient from two biological replicates. All selection coefficient values are shown regardless of whether they are significantly different than 0. Mutations with no fitness measurement in either replicate are shown in grey. Regions corresponding to the signal sequence (yellow), α-helices (green), β-strands (orange), the Ω-loop (white) and key active site residues (*) are shown above the heat map. Positions with a statistically significant difference in selection coefficient compared to 37°C in LB media can be found in **Fig S6**.

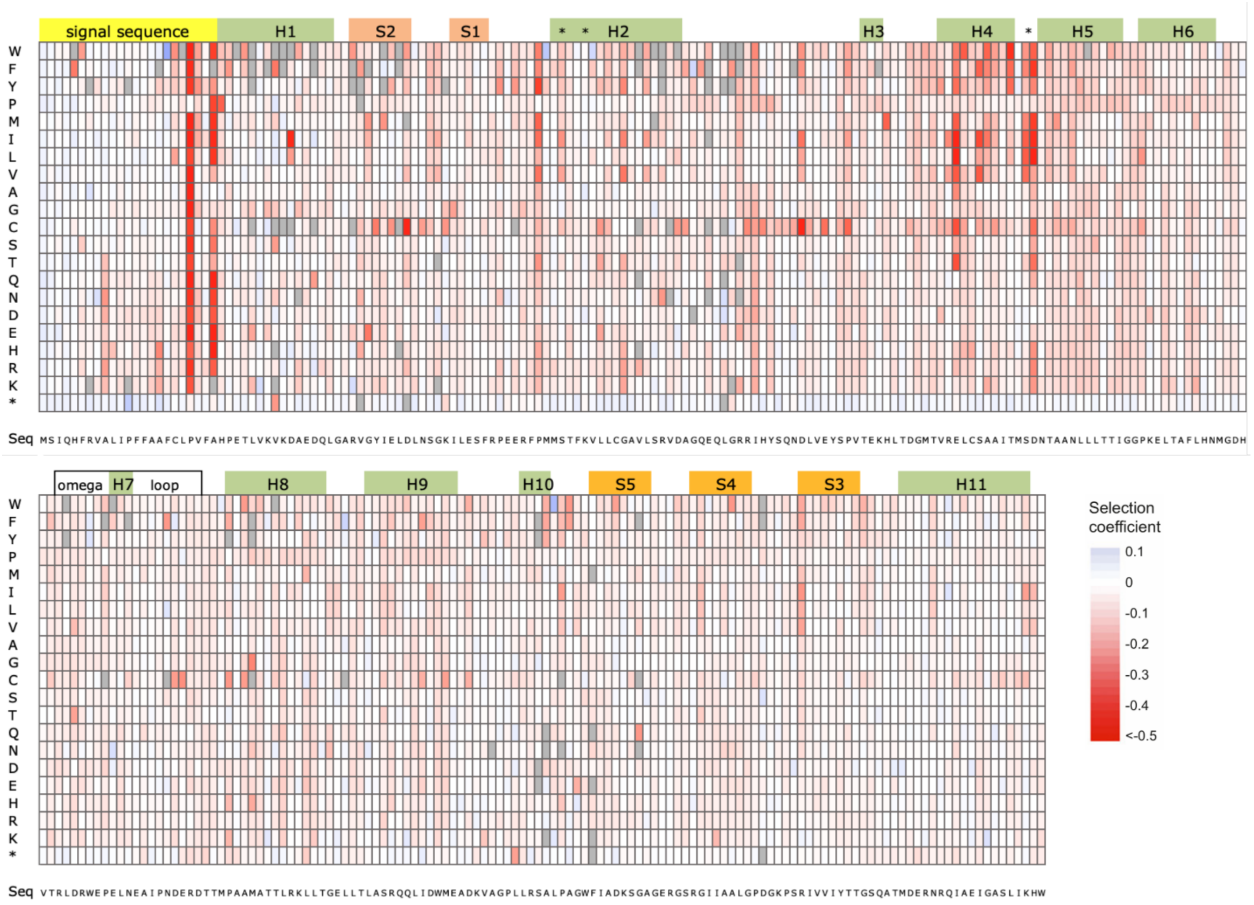


**Figure S4. Collateral fitness effects of mutations in TEM-1 at 42°C in LB media**. Heat map shows the weighted mean selection coefficient from two biological replicates. All selection coefficient values are shown regardless of whether they are significantly different than 0. Mutations with no fitness measurement in either replicate are shown in grey. Regions corresponding to the signal sequence (yellow), α-helices (green), β-strands (orange), the Ω-loop (white) and key active site residues (*) are shown above the heat map. Positions with a statistically significant difference in selection coefficient compared to 37°C in LB media can be found in **Fig S7-8**.

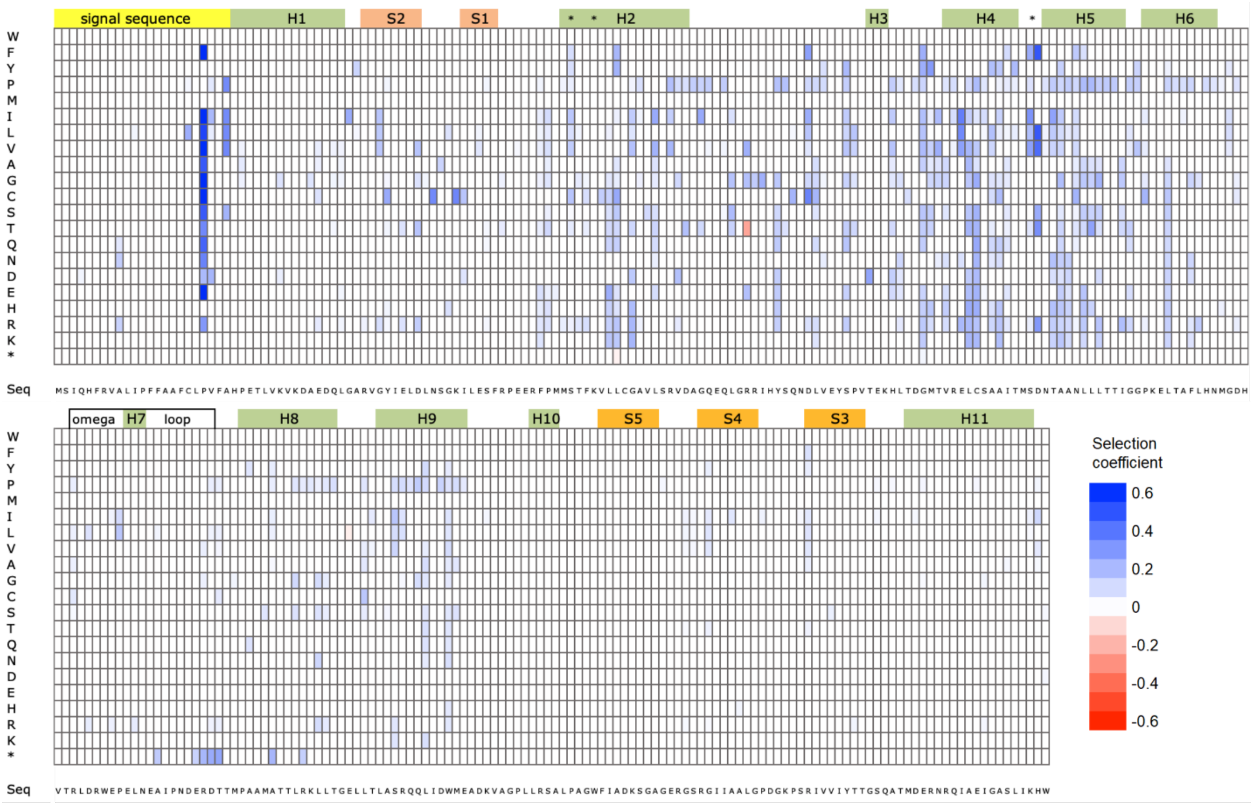


**Figure  S5. Significant changes in selection coefficients upon switching from 37°C to 30°C in LB media as analyzed by Student's t-test using data from individual codons.** Heat map shows statistically significant differences ($P < 0.01$). Blue indicates mutations less deleterious at 30°C, red indicates mutations more deleterious at 30°C. Regions corresponding to the signal sequence (yellow), α-helices (green), β-strands (orange), the Ω-loop region and key active site residues (*) are shown above the heat map.

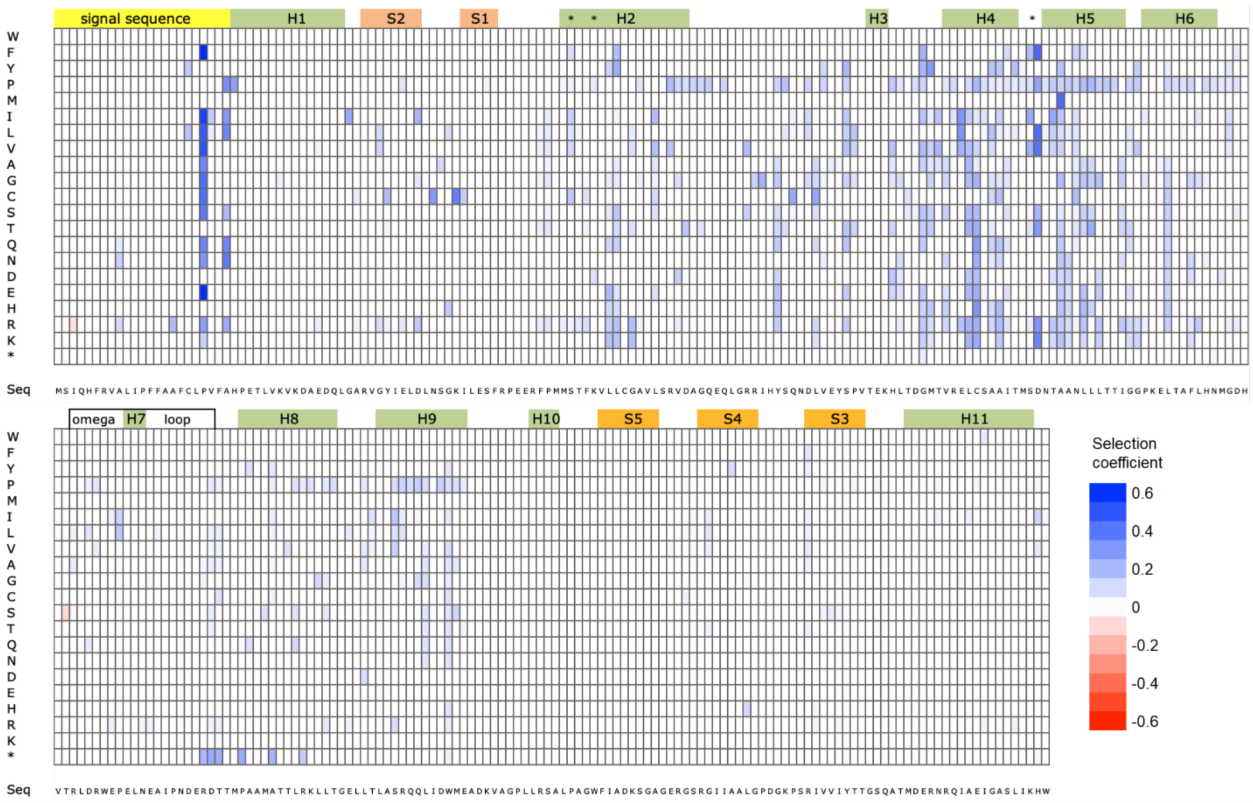


**Figure S6. Significant changes in selection coefficients upon switching from LB media to M9 minimal media at 37°C as analyzed by Student's t-test using data from individual codons**. Heat map shows statistically significant differences (*P* < 0.01). Blue indicates mutations less deleterious in M9 media, red indicates mutations more deleterious in M9 media. Regions corresponding to the signal sequence (yellow), α-helices (green), β-strands (orange), the Ω loop region and key active site residues (*) are shown above the heat map.

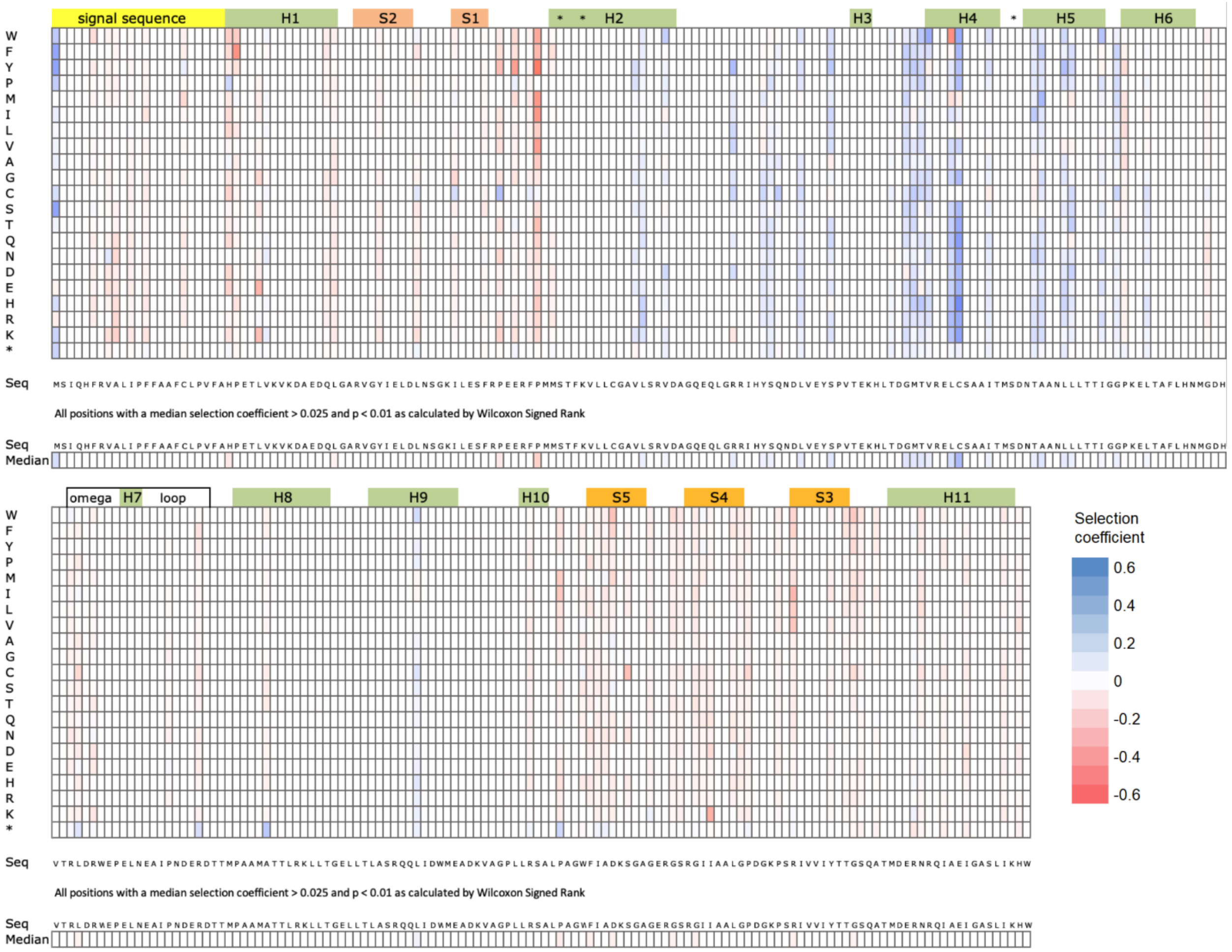


**Figure S7. Significant changes in selection coefficients upon switching from 37°C to 42°C in LB media as analyzed by the Wilcoxon signed rank test analysis**. Heat map shows statistically significant differences ($P < 0.01$). Blue indicates positions with mutations less deleterious at 42°C, red indicates positions with mutations more deleterious at 42°C. Median selection coefficients for each position are shown in the row labeled "Median." Regions corresponding to the signal sequence (yellow), α-helices (green), β-strands (orange), the Ω loop region and key active site residues (*) are shown above the heat map.

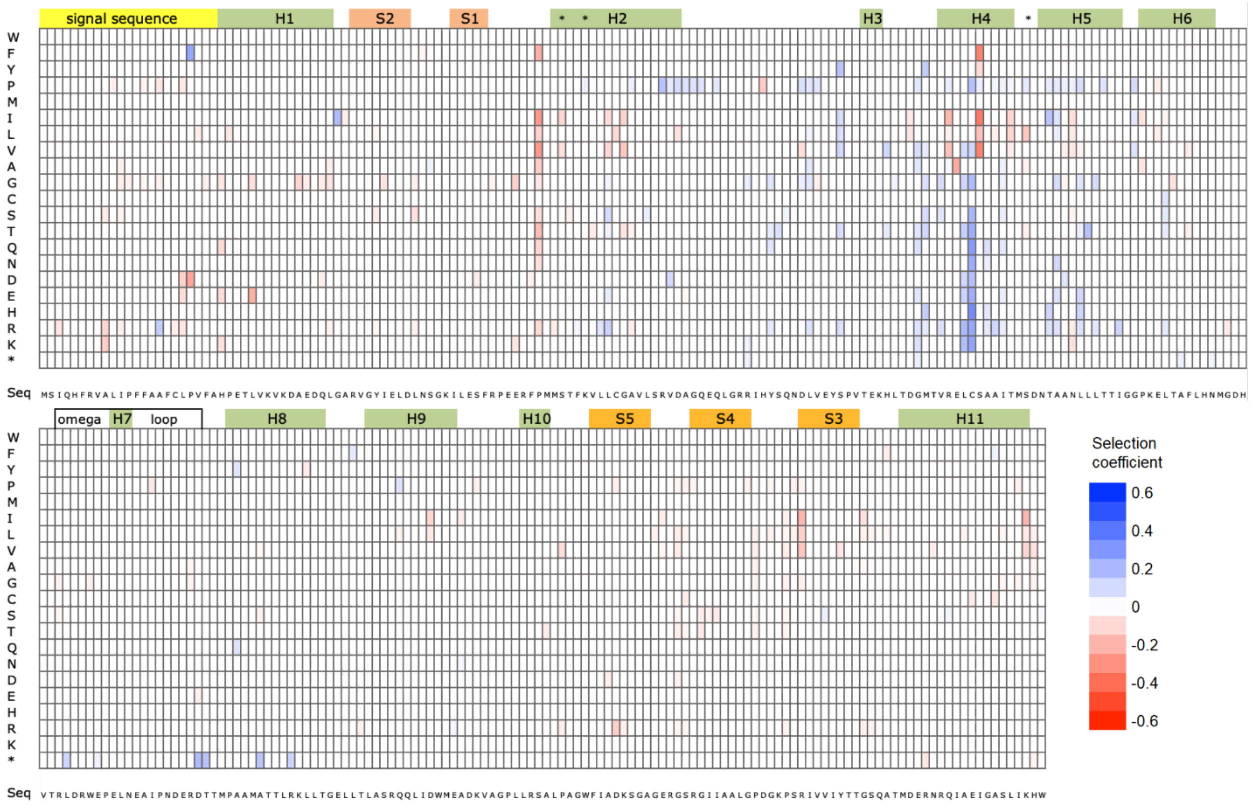


**Figure S8. Significant changes in selection coefficients upon switching from 37°C to 42°C in LB media as analyzed by Student's t-test using data from individual codons.** Heat map shows statistically significant differences ($P < 0.01$) in selection coefficients. Blue indicates mutations less deleterious at 42°C, red indicates mutations more deleterious at 42°C. Regions corresponding to the signal sequence (yellow), α-helices (green), β-strands (orange), the Ω loop region and key active site residues (*) are shown above the heat map.

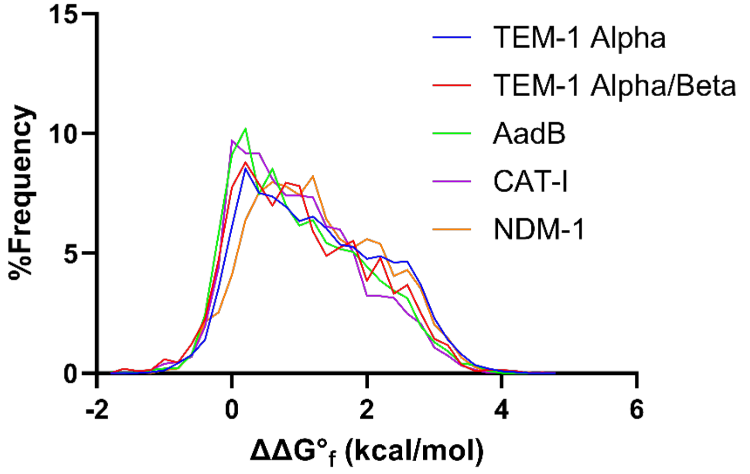


**Figure S9** Comparison of the distribution of ΔΔG°$_f$ for amino acid substitutions predicted by ThermoMPNN for the alpha (blue) and alpha/beta (red) domains of TEM-1 and AadB (green), CAT-I (purple), and NDM-1 (orange).

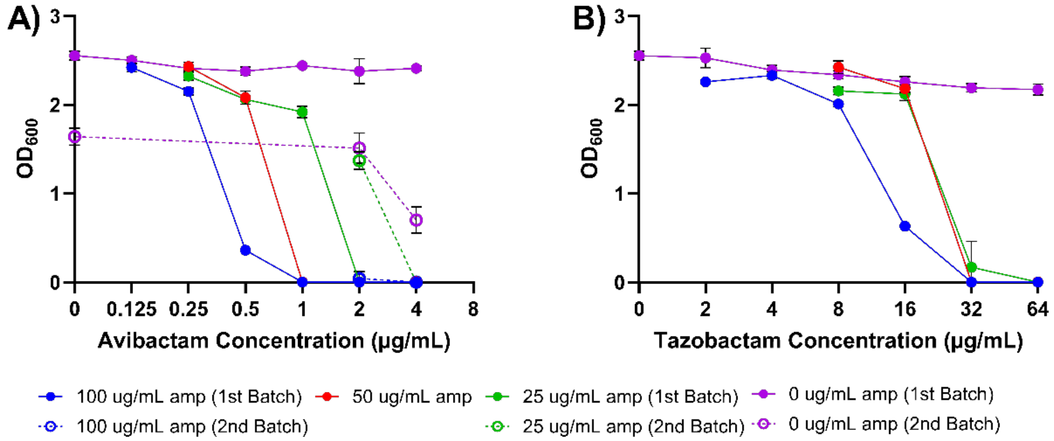


**Figure S10.** Minimum inhibitory concentration (MIC) for ampicillin (Amp) of NEB5-alpha F'Iq cells expressing TEM-1 in the presence of **A**) Avi and **B**) Tazo. Growth was measured as the OD600 after overnight growth at 37°C. Error bars indicate the standard deviation of the measurements which were calculated from biological replicates ($n$ = 3). Color indicates the concentration of Amp: blue (100 μg/mL), red (50 μg/mL), green (25 μg/mL), and purple (0 μg/mL). Solid lines indicate measurements from the first batch of Avi and Tazo. Dotted lines indicate measurements from the second batch of Avi.

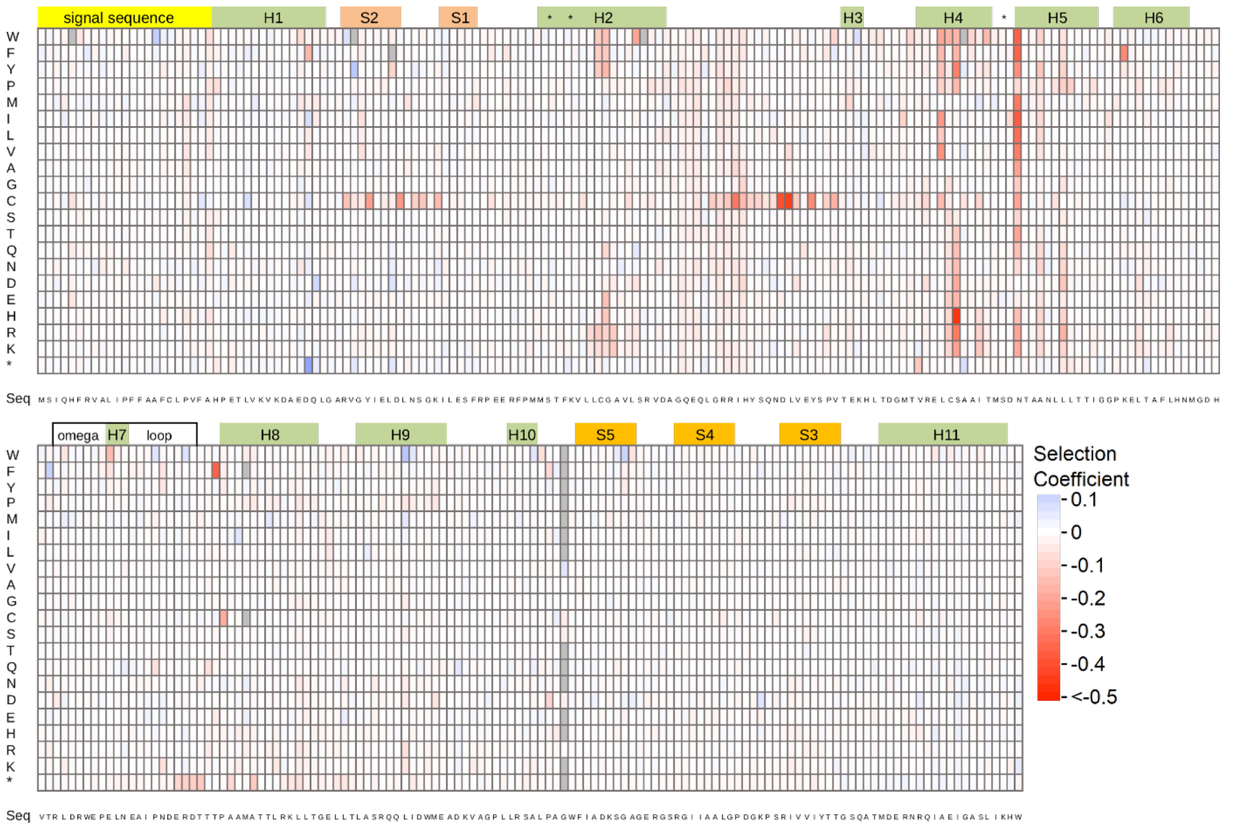


**Figure S11. Collateral fitness effects of mutations in TEM-1(M182T) at 37°C in LB media.** Weighted mean selection coefficients of two biological replicates are shown. These selection coefficients reflect the mean growth rates of mutants over ten generations post-TEM-1(M182T) induction in LB media relative to cells expressing TEM-1(M182T). Gray cells indicate mutants for which fitness was not measured in at least 1 replicate of DMS. Regions corresponding to the signal sequence (yellow), alpha-helices (green), beta-strands (orange), the Ω-loop (white) and three key active site residues (*) are shown.

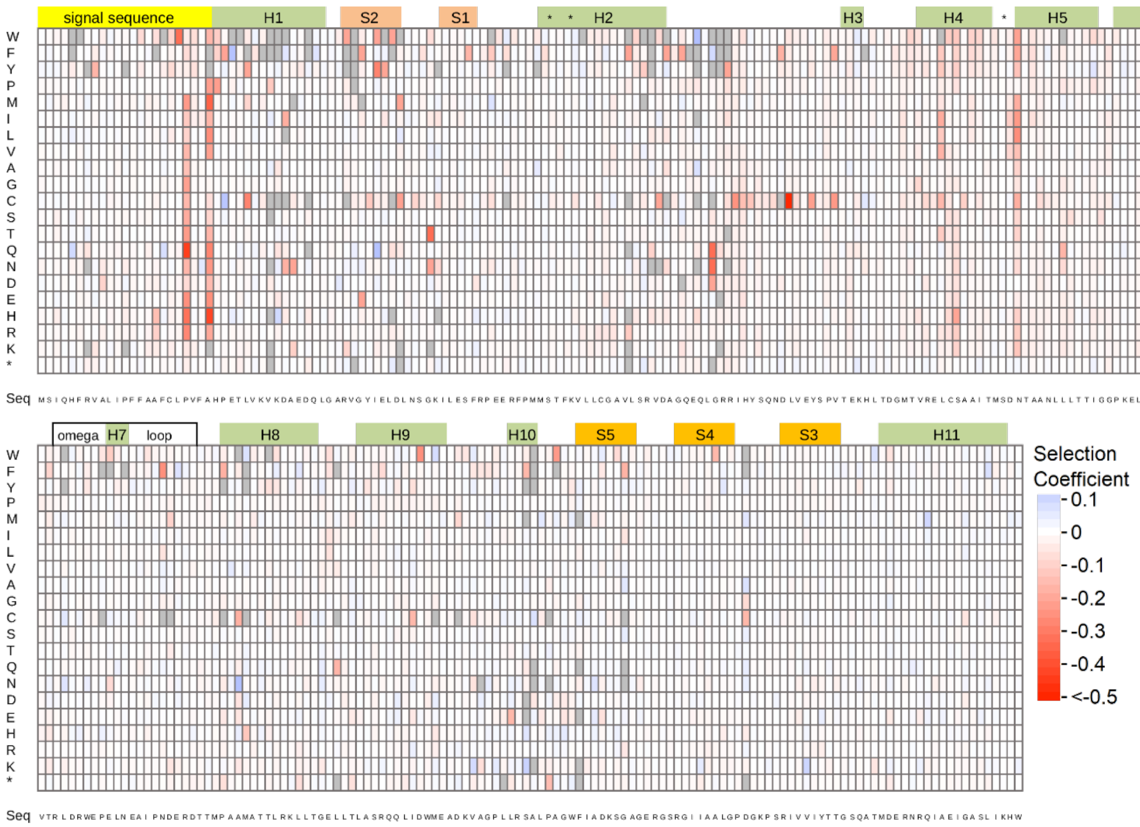


**Figure S12. Collateral fitness effects of mutations in TEM-1 at 37°C in LB media supplemented with Avi.** Weighted mean selection coefficients of two biological replicates are shown. These selection coefficients reflect the mean growth rates of mutants over ten generations post-TEM-1 induction in LB media supplemented with 2 μg/mL Avi relative to cells lacking a TEM-1 mutation. Gray cells indicate mutants for which fitness was not measured in at least 1 replicate of DMS. Regions corresponding to the signal sequence (yellow), alpha-helices (green), beta-strands (orange), the Ω-loop (white) and three key active site residues (*) are shown.

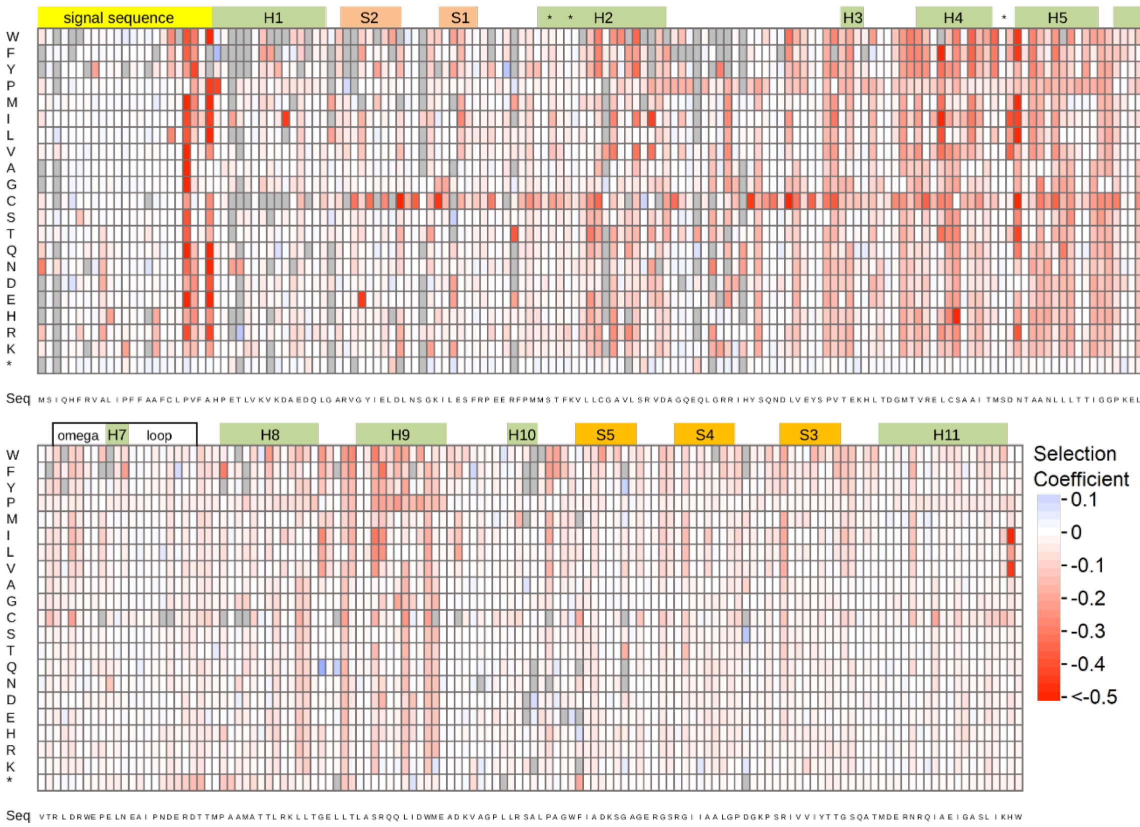


**Figure S13. Collateral fitness effects of mutations in TEM-1 at 37°C in LB media supplemented with Tazo.** Weighted mean selection coefficients of two biological replicates are shown. These selection coefficients reflect the mean growth rates of mutants over ten generations post-TEM-1 induction in LB media supplemented with 64 µg/mL Tazo relative to cells lacking a TEM-1 mutation. Gray cells indicate mutants for which fitness was not measured in at least 1 replicate of DMS. Regions corresponding to the signal sequence (yellow), alpha-helices (green), beta-strands (orange), the Ω-loop (white) and three key active site residues (*) are shown.

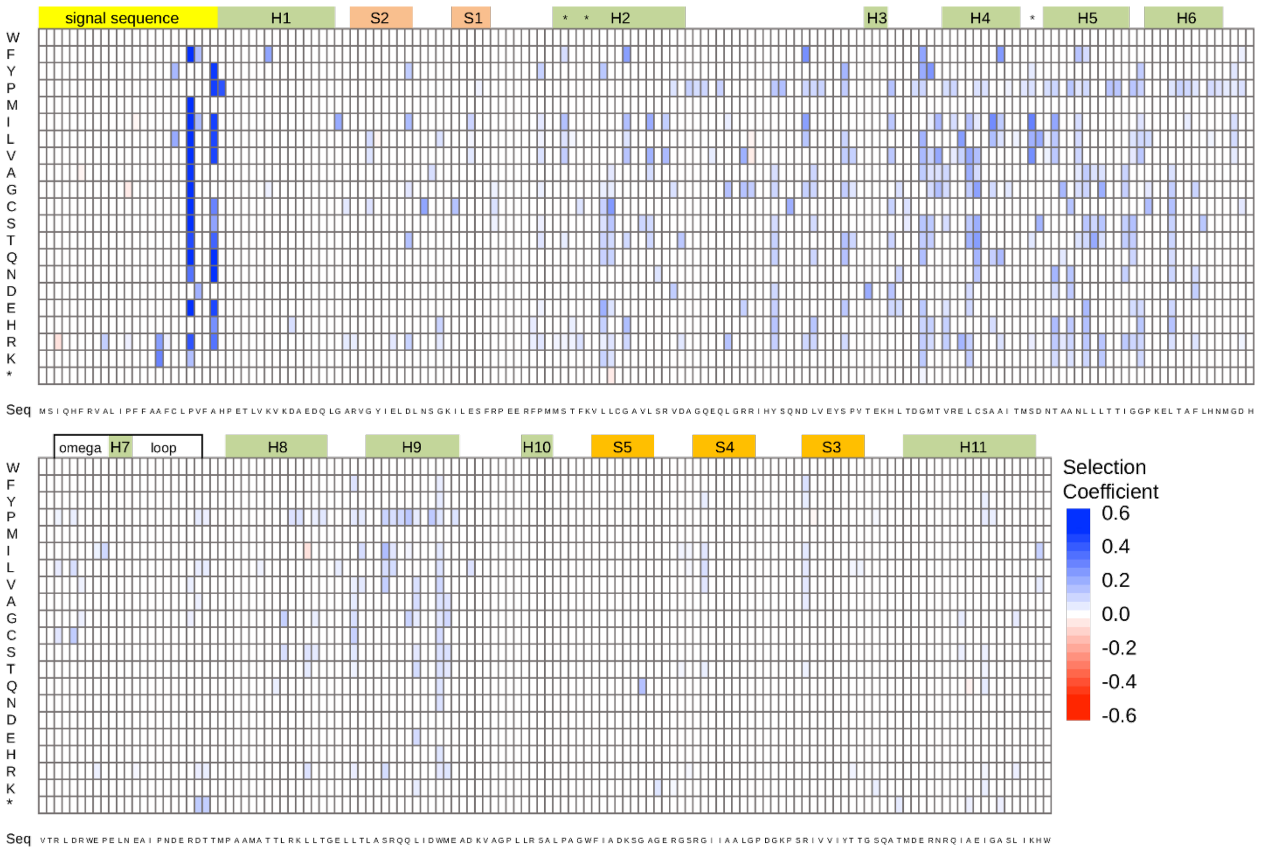


**Figure S14. Significant changes in selection coefficients in LB media at 37°C upon adding the M182T mutation as analyzed by Student's t-test using data from individual codons.** Heat map shows statistically significant differences ($P < 0.01$) in selection coefficients. Blue indicates mutations less deleterious with M182T and red indicates mutations more deleterious with M182T. Regions corresponding to the signal sequence (yellow), alpha-helices (green), beta-strands (orange), the *Ω* loop region and key active site residues (*) are shown above the heat map.

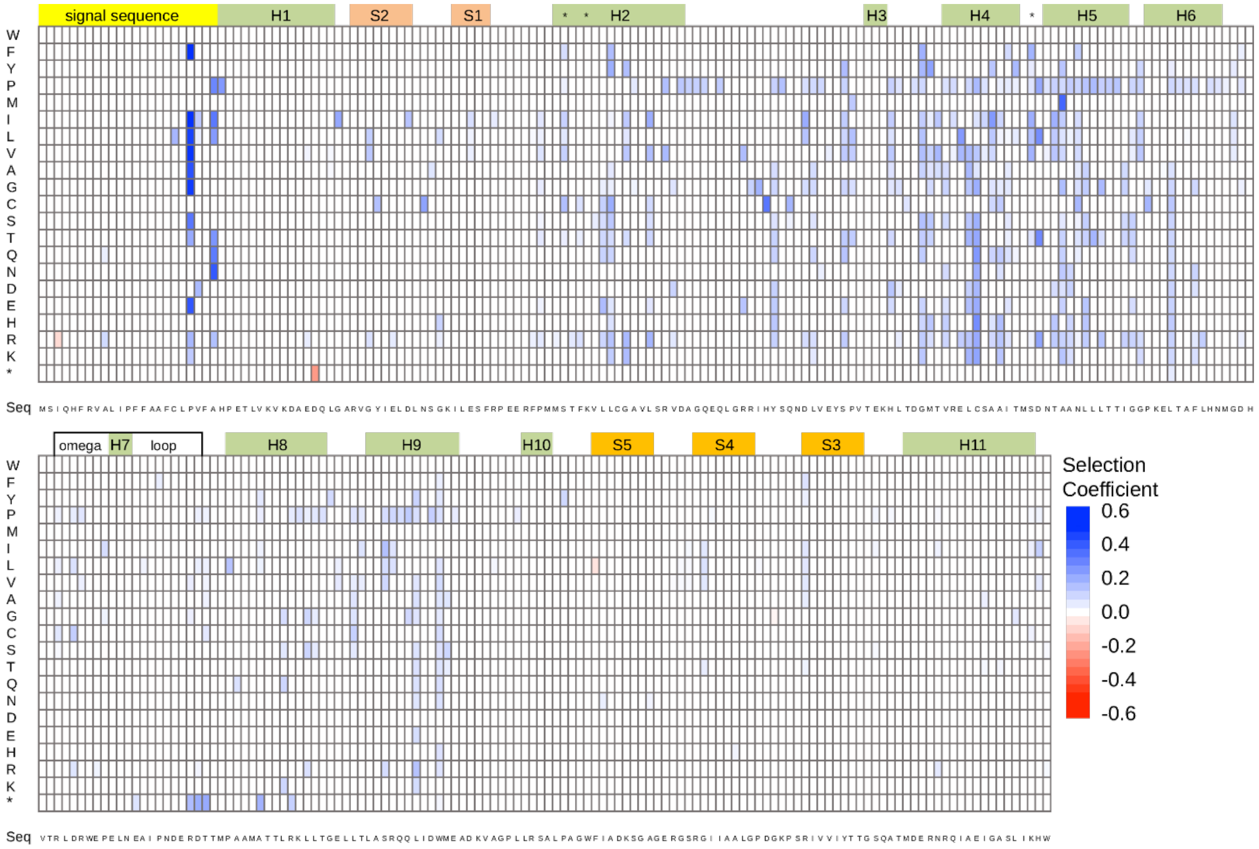


**Figure S15. Significant changes in selection coefficients at 37°C when LB media is supplemented with Avi as analyzed by Student's t-test using data from individual codons.** Heat map shows statistically significant differences ($P < 0.01$) in selection coefficients. Blue indicates mutations less deleterious with Avi and red indicates mutations more deleterious with Avi. Regions corresponding to the signal sequence (yellow), alpha-helices (green), beta-strands (orange), the *Ω* loop region and key active site residues (*) are shown above the heat map.

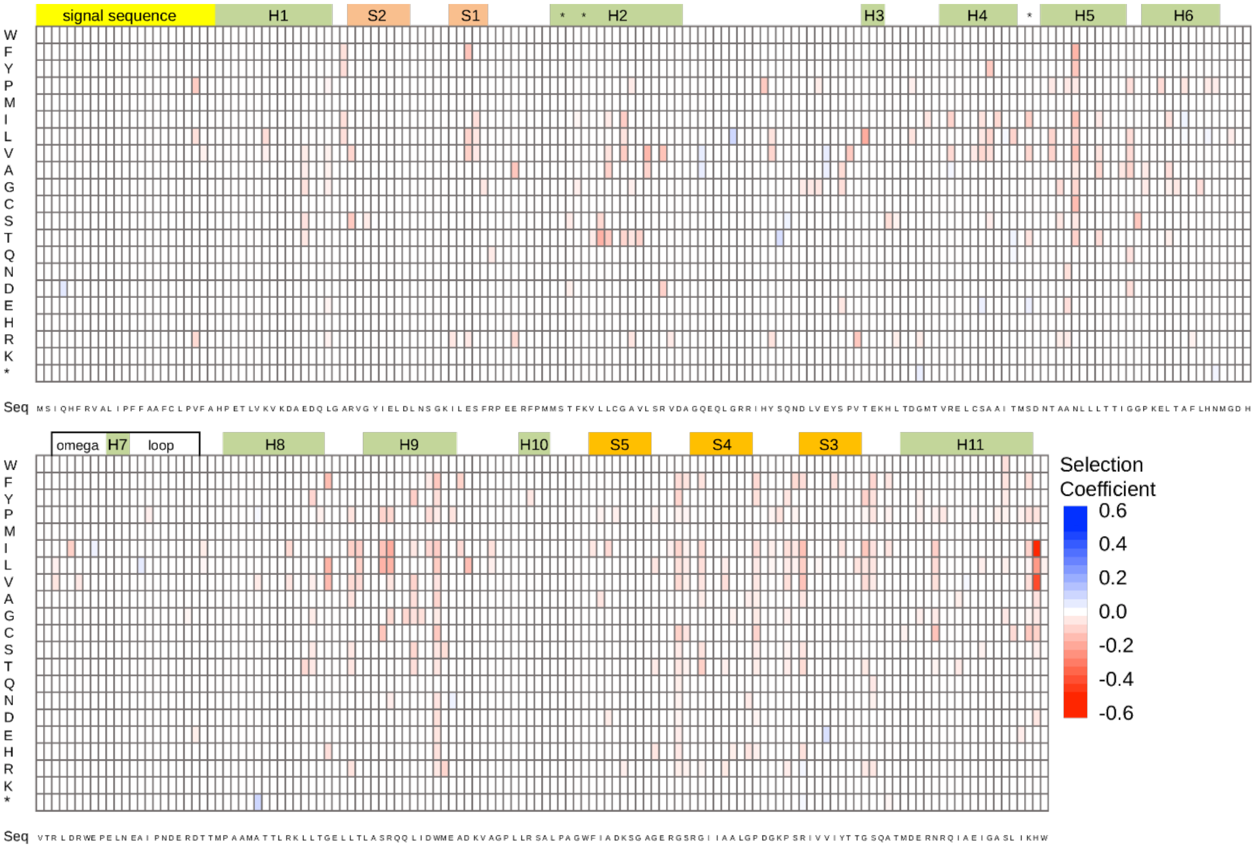


**Figure S16. Significant changes in selection coefficients at 37°C when LB media is supplemented with Tazo as analyzed by Student's t-test using data from individual codons.** Heat map shows statistically significant differences ($P < 0.01$) in selection coefficients. Blue indicates mutations less deleterious with Tazo, red indicates mutations more deleterious with Tazo. Regions corresponding to the signal sequence (yellow), alpha-helices (green), beta-strands (orange), the $\Omega$ loop region and key active site residues (*) are shown above the heat map.

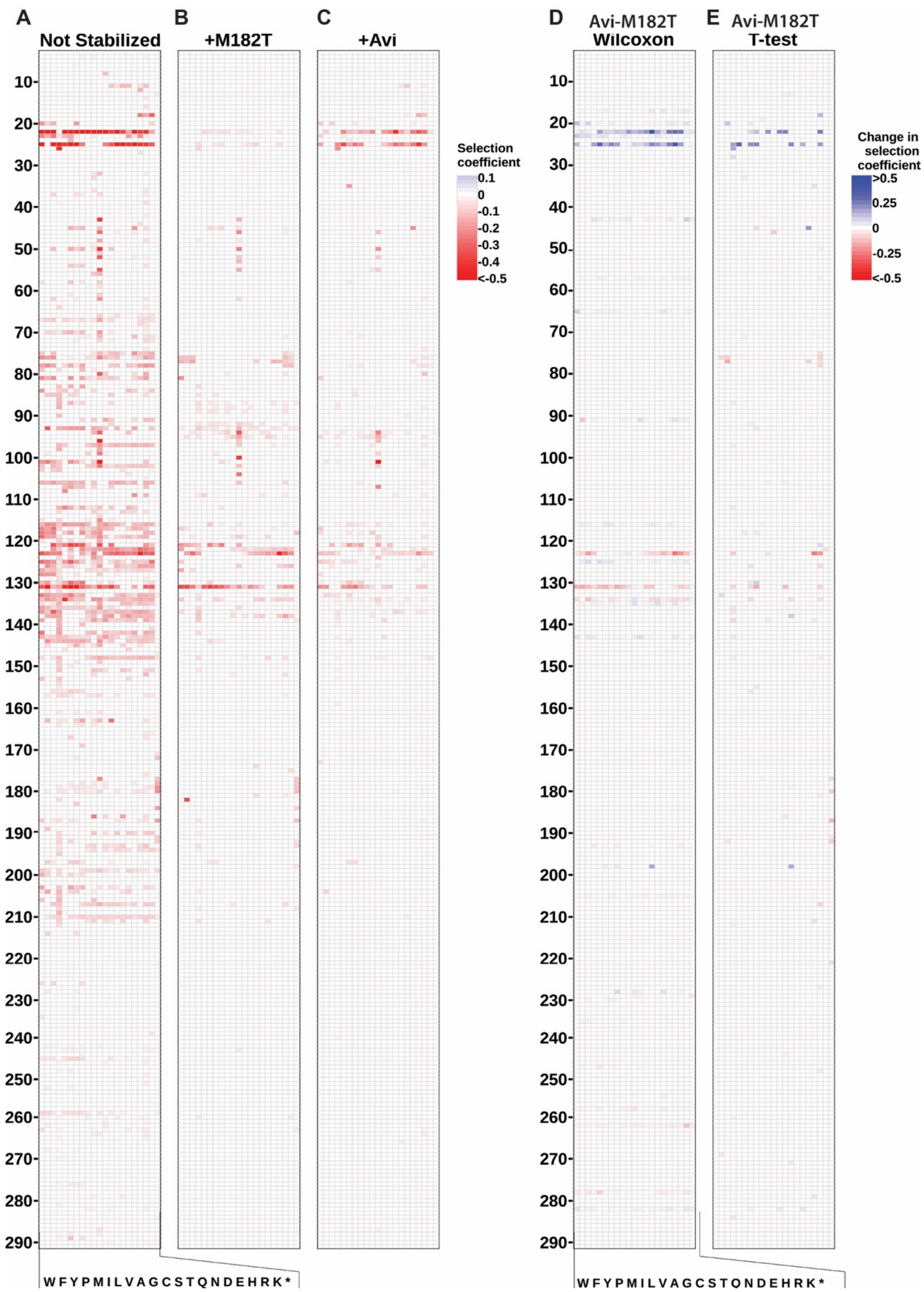


**Figure S17.** Differences in rescue of deleterious CFEs by M182T and Avi. Heat map of weighted mean selection coefficients for mutations that caused fitness effects ($P < 0.01$ in both replica experiments) under three different circumstances: **A**) TEM-1 **B**) TEM-1 with M182T **C**) TEM-1 in presence of Avi. Each column represents the amino acid substitution (indicated by single-letter codes at the bottom), and each row represents a position in the protein sequence (numbered on the left). Selection coefficients are relative to the wildtype under those conditions and color-coded according to the scale on the right, with blue indicating positive effects, white indicating neutral effects, and red indicating deleterious effects. Significant changes in selection coefficient upon shift from Avi to M182T as evaluated by **D**) Wilcoxon signed rank analysis and **E**) Student's t-test using data from individual codons.

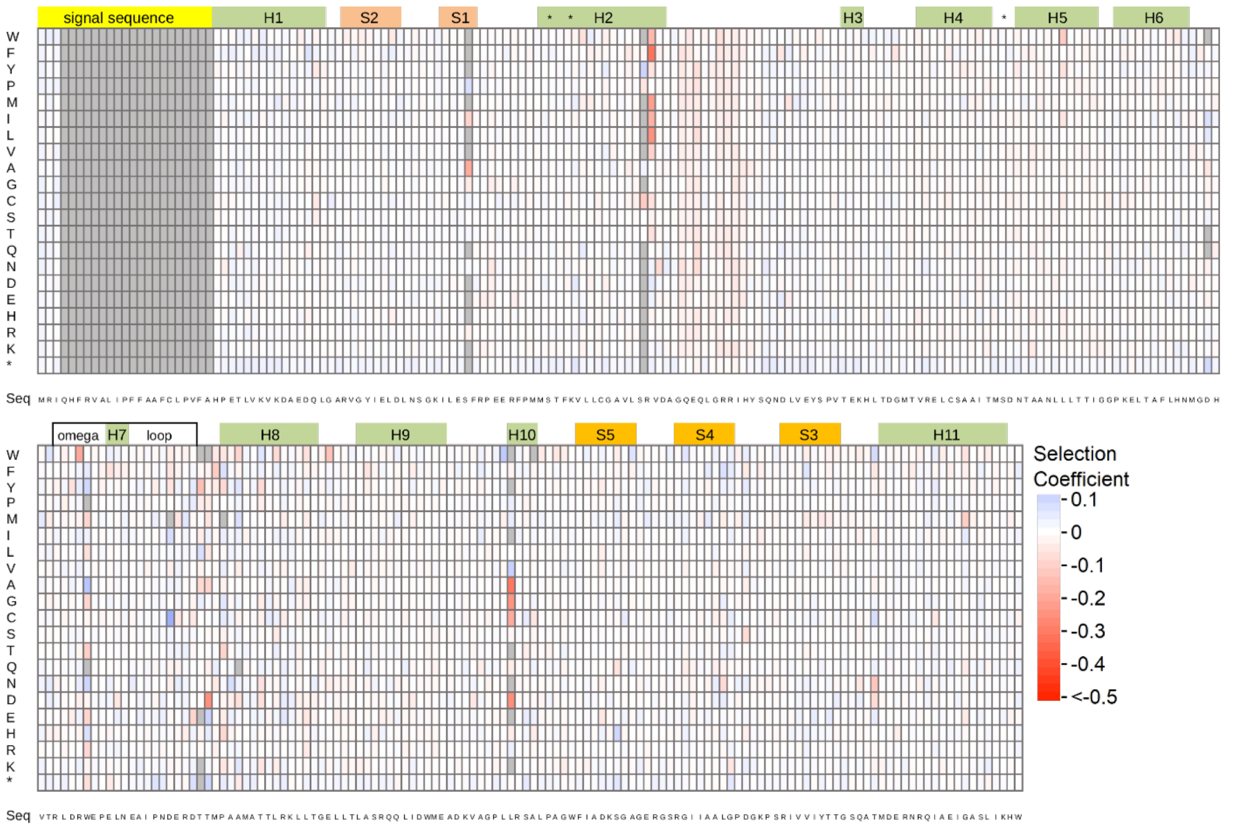


**Figure S18 Collateral fitness effects of mutations in ΔssTEM-1 at 37°C in LB media.** Heat map shows the weighted mean selection coefficient from two biological replicates. All selection coefficient values are shown regardless of whether they are significantly different than 0. Gray cells outside the signal sequence indicate mutants for which fitness was not measured in at least 1 replicate of DMS. Regions corresponding to the signal sequence (yellow), alpha-helices (green), beta-strands (orange), the Ω-loop (white) and three key active site residues (*) are shown.

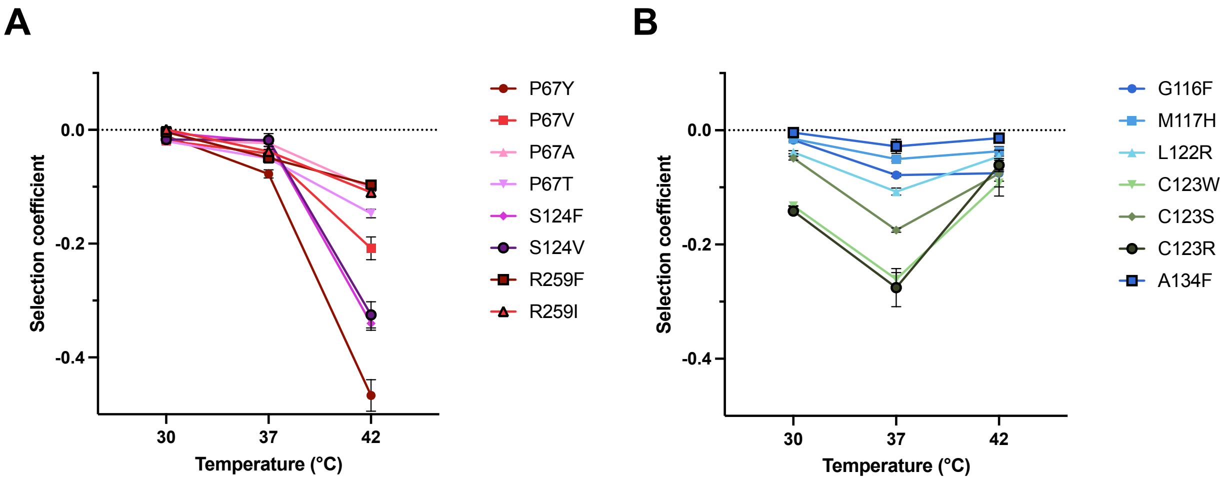


**Figure S19.** Temperature-dependent selection coefficients of TEM-1 mutants in LB media as measured by monoculture growth experiments. Error bars represent standard deviation from biological replicates (*n* ≥ 3). **(A)** mutations that are most deleterious at 42°C. **(B)** mutations that are most deleterious at 37°C. All selection coefficients are relative to WT TEM-1 at the same temperature. Data tabulated in **Table S2**.

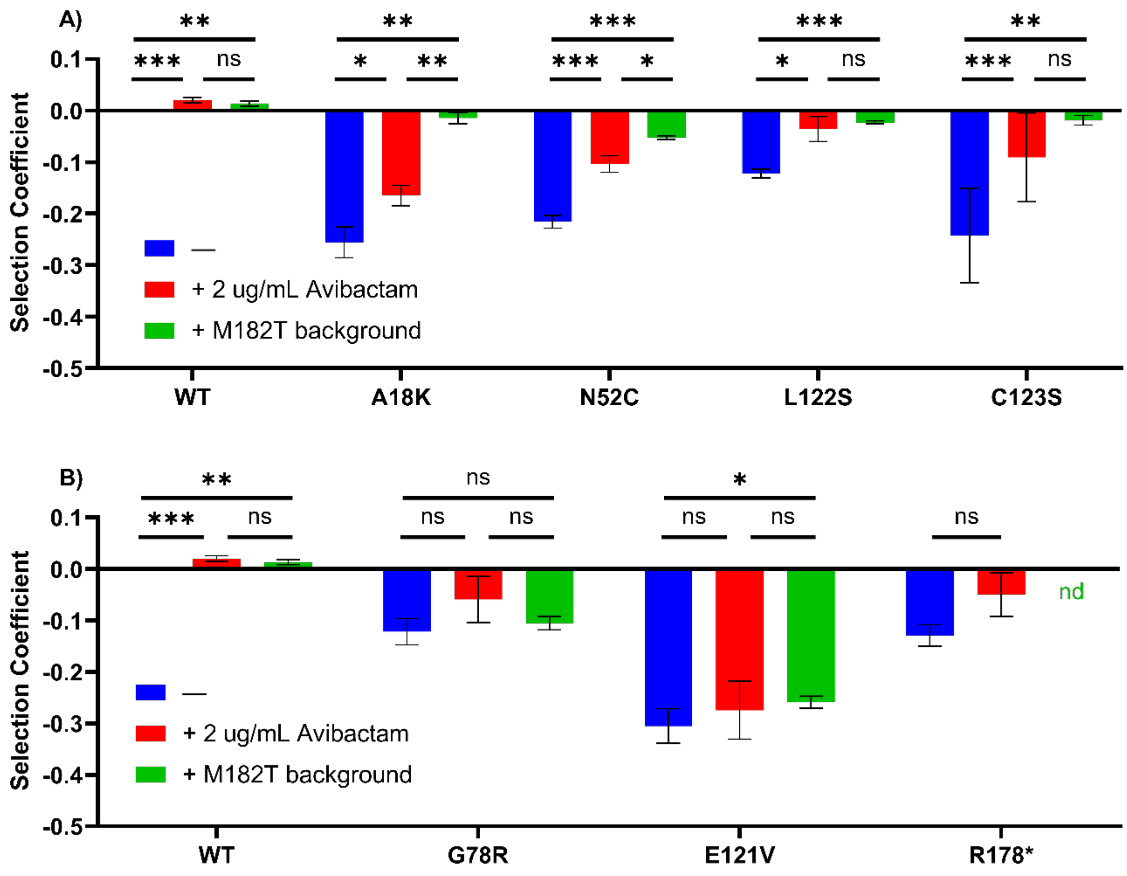


**Figure S20 A**) Significant and **B**) non-significant recovery of deleterious collateral fitness effects of select mutants by Avi and M182T as measured by monoculture growth experiments in LB media at 37°C. Blue, without Avi or M182T; red, with Avi; green, with M182T; nd = not determined. Error bars indicate the standard deviation from biological replicates ($n \geq 3$). $P$-values of t-tests: ns, $P \geq 0.05$; *, $P < 0.05$; **, $P < 0.01$; ***, $P < 0.001$. Cells were grown in LB media at 37°C. Data tabulated in **Table S3**.

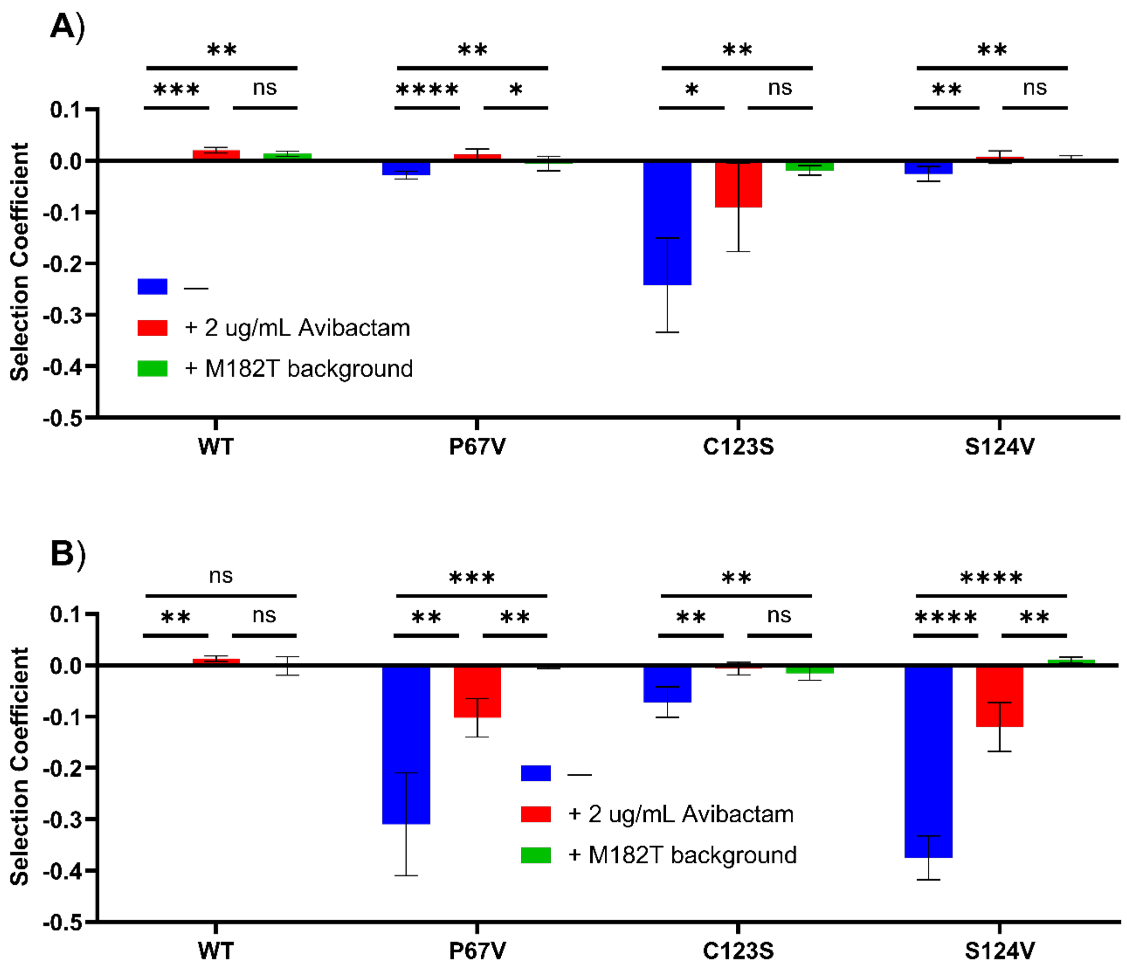


**Figure S21** Recovery of collateral fitness effects of select mutants with Avi and M182T in LB media at A) 37°C and B) 42°C as measured by monoculture growth experiments. All selection coefficients are relative to WT TEM-1 without M182T or Avi at the same temperature. Blue, without Avi or M182T; red, with Avi; green, with M182T. Error bars indicate the standard deviation from biological replicates ($n \geq 3$). *P*-values of t-tests: ns, $P \geq 0.05$; *, $P < 0.05$; **, $P < 0.01$; ***, $P < 0.001$; ****, $P < 0.0001$. Data tabulated in **Table S3.**

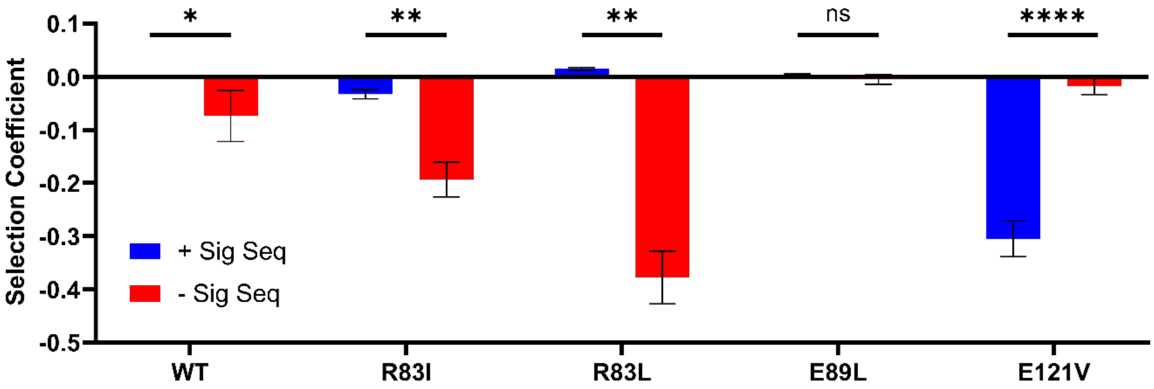


**Figure S22** Effect of the signal sequence on collateral fitness effects caused by select mutants as measured by monoculture growth experiments. All values are relative to WT TEM-1 with signal sequence. Blue, TEM-1 with signal sequence; red, ΔssTEM-1. Error bars indicate the standard deviation from biological replicates ($n \geq 3$). $P$-values of t-tests: ns, $P \geq 0.05$; *, $P < 0.05$; **, $P < 0.01$; ***, $P < 0.001$; ****, $P < 0.0001$. Cells were grown in LB media at 37°C. Data tabulated in **Table S4.**

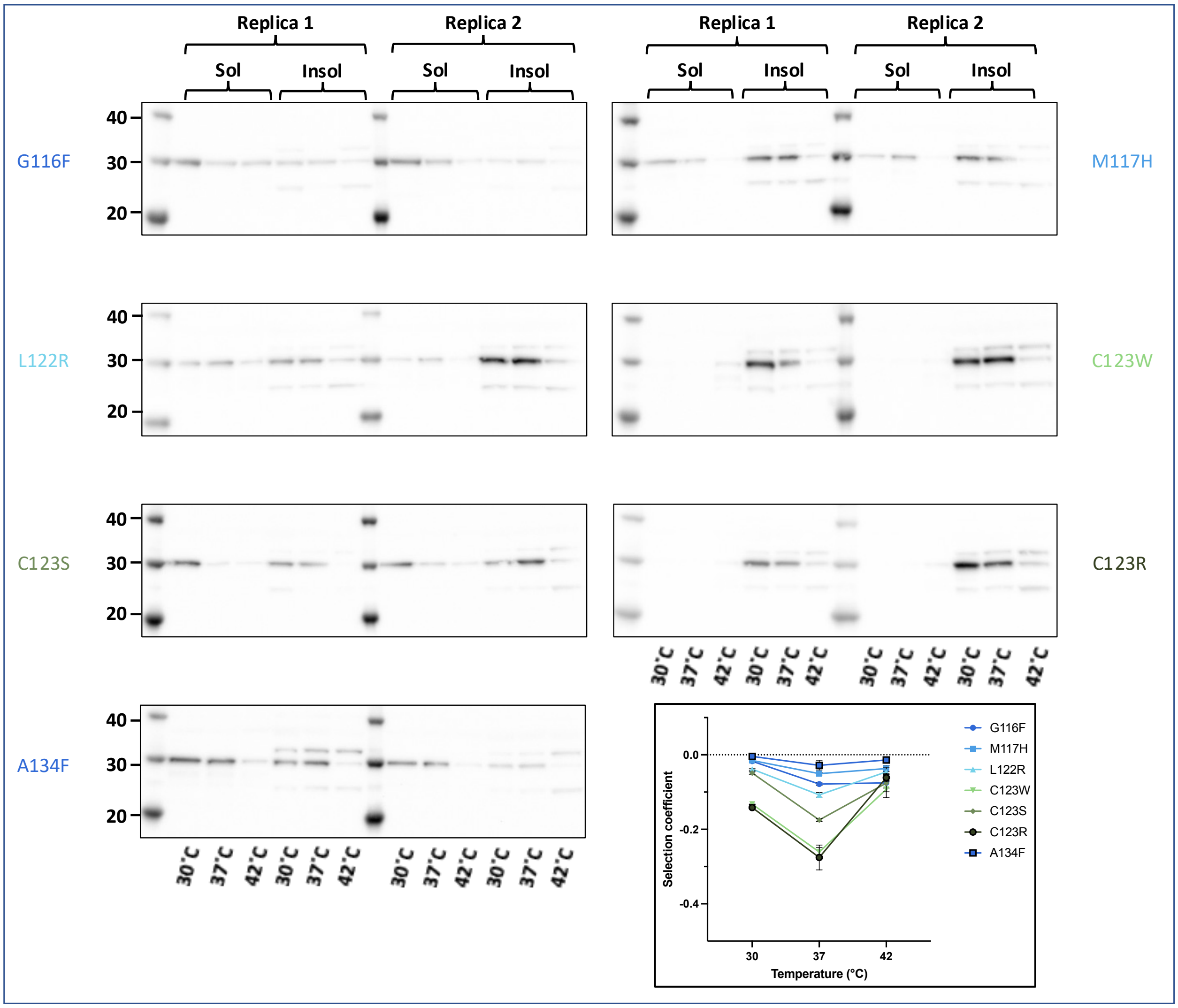


**Figure S23.** Western blots with TEM-1 antisera of the soluble and insoluble fractions of cells expressing TEM-1 mutants that are most deleterious at 37°C. For each variant, soluble and insoluble fractions from cells grown at 30°C, 37°C, and 42°C are arranged from left to right as indicated by the key at the top and bottom. Two biological replicates are shown. Molecular weight markers (in kDa) are indicated on the left. The matureTEM-1 is the prominent band at 30 kDa and preTEM-1 is the next highest band (when present). The temperature dependence of the selection coefficients for all mutants are included for reference (bottom right).

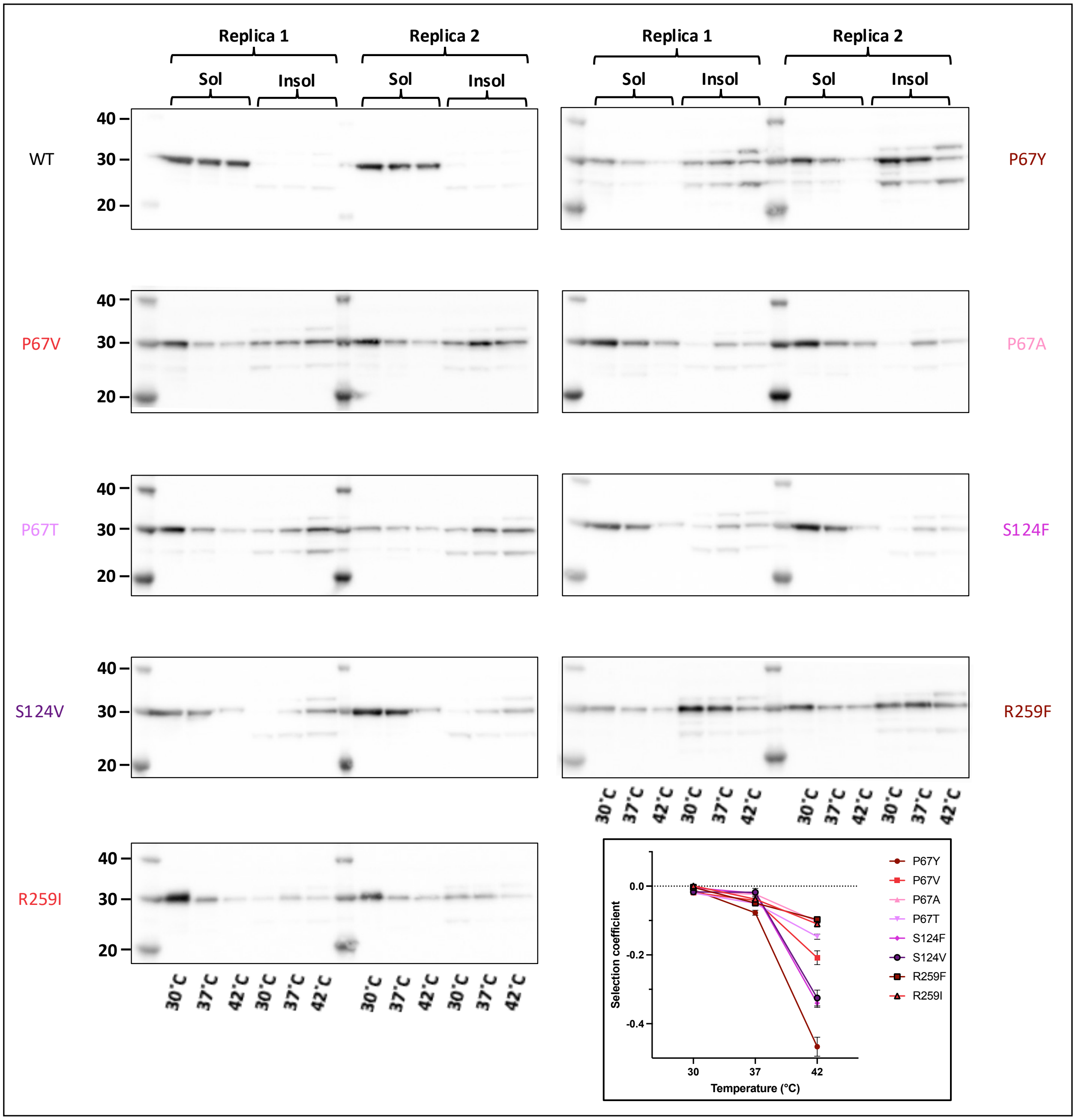


**Figure S24.** Western blots with TEM-1 antisera of the soluble and insoluble fractions of cells expressing TEM-1 mutants that are most deleterious at 42°C. For each variant, soluble and insoluble fractions from cells grown at 30°C, 37°C, and 42°C are arranged from left to right as indicated by the key at the top and bottom. Two biological replicates are shown. Molecular weight markers (in kDa) are indicated on the left. The matureTEM-1 is the prominent band at 30 kDa and preTEM-1 is the next highest band (when present). The temperature dependence of the selection coefficients for all mutants are included for reference (bottom right).

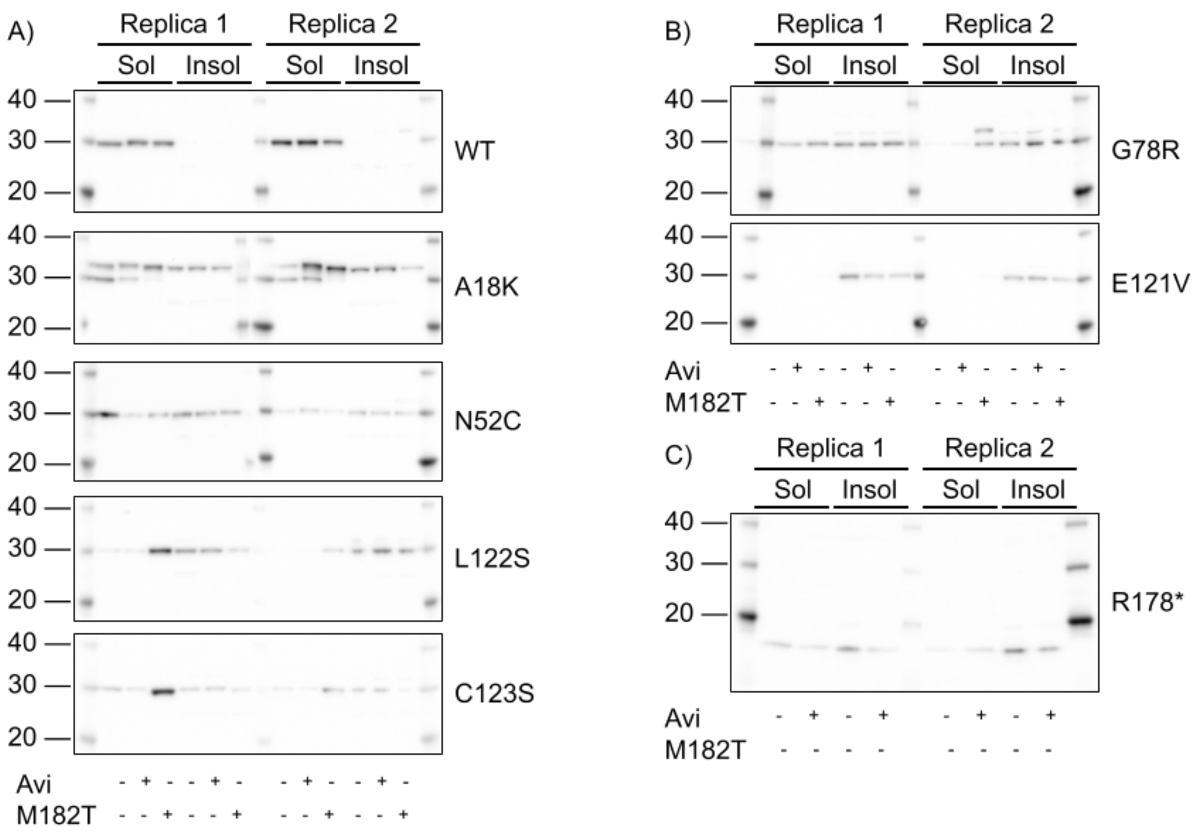


**Figure S25** Western blots with TEM-1 antisera of the soluble and insoluble fractions of cells expressing TEM-1 mutants from cells grown in the presence of Avi, in the presence of the M182T mutation, or with neither. Two biological replicates are shown. Molecular weight markers (in kDa) are indicated on the left. The matureTEM-1 is the prominent band at 30 kDa and preTEM-1 is the next highest band (when present). **A**) Mutants that demonstrated a large magnitude change in selection coefficient in the presence of Avi or M182T. WT = no mutations. **B**) Mutants that did not demonstrate a large magnitude change in selection coefficient in the presence of Avi or M182T. **C**) The R178* mutant.

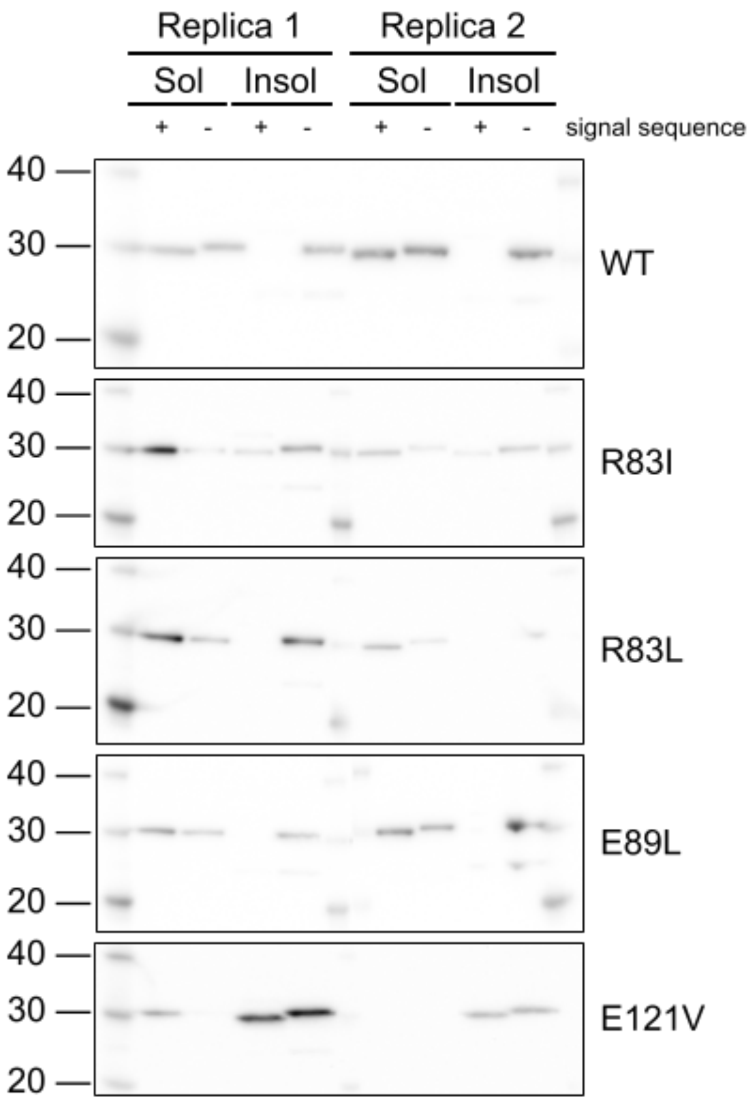


**Figure S26.** Western blots with TEM-1 antisera of the soluble and insoluble fractions of cells expressing TEM-1 mutants in the presence or absence of the signal sequence. Two biological replicates are shown. Molecular weight markers (in kDa) are indicated on the left. The mature TEM-1 is the prominent band at 30 kDa (preTEM-1 was not observed in these experiments).

**Table S1.** Statistics on CFEs as measure by deep mutational scanning

| Temp. (°C) | Media | Addition/mutation | Percent deleterious[a] | Selection coefficient | |
|---|---|---|---|---|---|
| | | | | Mean | Median |
| 30 | LB | - | 3.7% | -0.0020 | +0.0012 |
| 37 | M9 | - | 6.3% | -0.0057 | +0.0020 |
| 37 | LB | - | 21.4% | -0.0345 | -0.0113 |
| 42 | LB | - | 32.0% | -0.0378 | -0.0243 |
| 37 | LB | 64 µg/ml tazobactam | 32.7% | -0.0477 | -0.0189 |
| 37 | LB | 2 µg/ml avibactam | 4.6% | -0.0070 | -0.0023 |
| 37 | LB | M182T | 5.0% | -0.0060 | -0.0015 |
| 37 | LB | Δ signal sequence | 1.4% | -0.0001 | +0.0012 |

[a] Percentage of mutations with deleterious effects (*P* <0.01 in both replica experiments)

**Table S2.** Selection coefficients at different temperatures as measured by monoculture growth experiments in LB media.

| Mutation | Selection coefficient[a] | | | Standard deviation[b] | | |
|---|---|---|---|---|---|---|
| | 30°C | 37°C | 42°C | 30°C | 37°C | 42°C |
| P67Y | -0.0108 | -0.0774 | -0.4670 | 0.0013 | 0.0071 | 0.0277 |
| P67V | -0.0188 | -0.0414 | -0.2081 | 0.0026 | 0.0054 | 0.0200 |
| P67A | -0.0163 | -0.0234 | -0.1033 | 0.0033 | 0.0054 | 0.0045 |
| P67T | -0.0197 | -0.0502 | -0.1469 | 0.0018 | 0.0044 | 0.0075 |
| G116F | -0.0171 | -0.0783 | -0.0752 | 0.0025 | 0.0033 | 0.0141 |
| M117H | -0.0153 | -0.0505 | -0.0369 | 0.0039 | 0.0037 | 0.0082 |
| L122R | -0.0385 | -0.1074 | -0.0457 | 0.0025 | 0.0062 | 0.0109 |
| C123R | -0.1415 | -0.2755 | -0.0611 | 0.0029 | 0.0333 | 0.0115 |
| C123S | -0.0488 | -0.1748 | -0.0758 | 0.0032 | 0.0036 | 0.0234 |
| C123W | -0.1326 | -0.2608 | -0.0924 | 0.0004 | 0.0116 | 0.0229 |
| S124F | -0.0048 | -0.0206 | -0.3402 | 0.0082 | 0.0032 | 0.0121 |
| S124V | -0.0168 | -0.0179 | -0.3251 | 0.0067 | 0.0113 | 0.0229 |
| A134F | -0.0040 | -0.0282 | -0.0139 | 0.0022 | 0.0123 | 0.0022 |
| R259F | -0.0039 | -0.0493 | -0.0970 | 0.0051 | 0.0031 | 0.0055 |
| R259I | 0.0007 | -0.0377 | -0.1099 | 0.0053 | 0.0037 | 0.0078 |

[a] Relative to WT TEM-1 without M182T or Avi at the same temperature.
[b] $n \geq 3$

**Table S3.** Selection coefficients in the presence and absence of Avi or M182T as measured by monoculture growth experiments in LB media.

| Fig. | Mutation | Temperature | Selection coefficient[a] | | | Standard deviation[b] | | |
|---|---|---|---|---|---|---|---|---|
| | | | – | Avi | M182T | – | Avi | M182T |
| S19 | none | 37 °C | 0.0000 | 0.0223 | 0.0136 | 0.0000 | 0.0023 | 0.0049 |
| | A18K | | -0.2556 | -0.1646 | -0.0145 | 0.0299 | 0.0197 | 0.0109 |
| | N52C | | -0.2157 | -0.1034 | -0.0524 | 0.0123 | 0.0160 | 0.0036 |
| | G78R | | -0.1219 | -0.0590 | -0.1053 | 0.0254 | 0.0445 | 0.0127 |
| | E121V | | -0.3052 | -0.2746 | -0.2587 | 0.0336 | 0.0562 | 0.0117 |
| | L122S | | -0.1221 | -0.0357 | -0.0227 | 0.0085 | 0.0243 | 0.0030 |
| | C123S | | -0.2048 | -0.0540 | -0.0187 | 0.0428 | 0.0321 | 0.0092 |
| | R178* | | -0.1292 | -0.0497 | n.d.[c] | 0.0208 | 0.0424 | n.d.[c] |
| S20 | none | | 0.0000 | 0.0203 | 0.0136 | 0.0000 | 0.0055 | 0.0049 |
| | P67V | | -0.0281 | 0.0121 | -0.0058 | 0.0076 | 0.0111 | 0.0138 |
| | C123S | | -0.2421 | -0.0905 | -0.0187 | 0.0915 | 0.0863 | 0.0092 |
| | S124V | | -0.0258 | 0.0068 | 0.0033 | 0.0144 | 0.0122 | 0.0067 |
| | none | 42 °C | 0.0000 | 0.0130 | -0.0013 | 0.0000 | 0.0053 | 0.0180 |
| | P67V | | -0.3098 | -0.1023 | -0.0032 | 0.1001 | 0.0376 | 0.0033 |
| | C123S | | -0.0719 | -0.0064 | -0.0156 | 0.0295 | 0.0126 | 0.0135 |
| | S124V | | -0.3751 | -0.1205 | 0.0100 | 0.0427 | 0.0474 | 0.0060 |

[a] Relative to WT TEM-1 without M182T or Avi at the same temperature.
[b] $n \geq 3$
[c] not determined

**Table S4.** Selection coefficients in the presence and absence of the signal sequence.

| Mutation | Selection coefficient[a] | | Standard deviation[b] | |
|---|---|---|---|---|
| | +ss | Δss | +ss | Δss |
| WT | 0.0000 | -0.0736 | 0.0000 | 0.0483 |
| R83L | 0.0148 | -0.3779 | 0.0030 | 0.0495 |
| R83I | -0.0328 | -0.1931 | 0.0089 | 0.0327 |
| E89L | 0.0032 | -0.0047 | 0.0023 | 0.0091 |
| E121V | -0.3052 | -0.0169 | 0.0336 | 0.0162 |

[a] Relative to WT TEM-1 with signal sequence.
[b] $n \geq 3$
